# High-Resolution Spatial Transcriptomics Reveals Fibroblast and Neuroimmune Microenvironments in Endometriosis Lesions

**DOI:** 10.1101/2025.08.20.671344

**Authors:** Caroline M. Haney, Elaheh Alizadeh, Meryl Sullivan, Joshua Lee, Jasmina Kuljancic, William F. Flynn, Paul Robson, Brian S. White, Danielle E. Luciano, Elise T. Courtois

## Abstract

Endometriosis is a chronic, systemic, inflammatory disease characterized by the presence of endometrium-like tissue growing outside of the uterus. One of its main symptoms is chronic pain and inflammation leading to a decreased quality of life. This is a common disease, as at least one in ten female-born individuals have endometriosis. Yet the understanding of the mechanisms that drive pain symptoms and disease progression remain poorly defined. This study establishes the precise spatial transcriptomic cartography of human ovarian and peritoneal lesions, two of the most commonly found lesions. We identified shared spatial features across lesion types, including immune cell infiltration, fibroblast specific compartments surrounding epithelial glands, and distinct distributions of neuronal and macrophage subsets.

We precisely defined sensory neuronal subtypes, and mapped their spatial location relative to immune cells. We further validate the epithelial-neuronal interactome, using an *in vitro* 3D model of peripheral sensory brain organoids co-cultured with human endometriosis epithelial and fibroblast cells. By mapping spatial cellular interactions and identifying conserved features across lesion types linked to pain, our study provides emerging insights into endometriosis pathophysiology, paving the way for the development of novel targeted therapeutic strategies.

## Introduction

Endometriosis is a chronic, systemic, inflammatory, and estrogen-dependent condition characterized by the growth of endometrium-like tissue outside of the uterus, termed lesions. These lesions are diverse and heterogeneous in morphology and are most commonly located on the pelvic peritoneum, ovaries, and invading surrounding organs. Endometriosis affects ∼7-10% of individuals assigned female at birth, with symptoms including heavy menstrual bleeding, chronic pelvic pain, infertility, and urinary or gastrointestinal dysfunction typically emerging during the reproductive years. However, diagnosis is often delayed by 7-10 years allowing the disease to progress and complications to accumulate, including the development of peripheral and central sensitization^1–6^. Endometriosis patients experience neuroplastic changes that amplify pain perception through heightened sensitivity of peripheral nociceptors and increased excitability of central pain pathways^6^. Endometriosis is therefore a lifelong disease with substantial morbidity, beginning in adolescence. Currently, there is no cure for endometriosis and available pharmacotherapies are limited to hormonal regulation and pain management.

Both of these treatment options are associated with significant side effects and limited efficacy and, most importantly, fail to treat the root cause of the disease and address the underlying pathophysiology^7^.

Histologically, endometriosis lesions are defined by the presence of endometrium-like glandular epithelial cells surrounded by fibroblasts, and an abundance of hemosiderin-laden macrophages. However, recent endometriosis single cell studies have expanded our understanding of the complex lesion microenvironment, revealing critical contributions from heterogeneous fibroblasts subpopulations, immune cells, and blood vessels^8–10^. Collectively, these studies suggest that endometriosis lesions drive immune cell recruitment and rely on a finely tuned balance of pro-inflammatory and immunosuppressive signals – mediated by diverse cell types including macrophages, dendritic cells (DCs), and others – to establish an immune-rich niche. Our previous work identified both shared and distinct cellular phenotypes between peritoneal and ovarian lesions, the two most common subtypes, underscoring the importance of resolving lesion heterogeneity to identify conserved therapeutic targets.

Local tissue remodeling and inflammation are central features of endometriosis-driven microenvironmental changes^8^. Recent studies have explored the role of neuro-immune signaling in the generation of endometriosis-associated pain, with a particular focus on macrophage-neuron cross-talk^11–19^. Macrophages secrete a range of pro-inflammatory cytokines, such as IL-1β, TNF-α, IL6 and prostaglandins, that can act on nearby nociceptive neurons to amplify pain signaling^12^. In addition, macrophages contribute to the growth of endometriosis lesions in mouse models by increasing angiogenesis, matrix remodeling, and tissue invasion^11^. Moreover, they secrete neurotrophic factors such as Brain-Derived Neurotrophic Factor (BDNF), Insulin Growth Factor-1 (IGF-1), and Nerve Growth Factor (NGF) which can stimulate neuronal sprouting^20^. Conversely, sensory neurons secrete neuropeptides such as Substance P (SP) and Calcitonin Gene-Related Peptide (CGRP) which signal back to macrophages and other immune cells to further enhance inflammation^21^. Recent work in a mouse model for endometriosis showed that blocking CGRP-RAMP1 signaling in macrophages reduced lesion size and pain symptoms^14^. Together, these findings suggest a model where reciprocal neuropeptide-receptor interactions lead to a feedforward loop of inflammation and pain.

However, translating these neuro-immune insights into human disease remains a major challenge. Mouse models, while useful, fail to fully recapitulate human lesions and their diversity. Additionally, immunohistochemistry-based studies have confirmed that nerve fibers are present near human endometriosis lesions in a broad sensory vs. sympathetic manner, but this approach lacks the resolution to define their molecular identity or spatial organization/interactions^11–13^. As a result, the neuronal landscape and full complexity of neuro-immune interactions in human endometriosis remains poorly understood. This is significant, as several studies have emphasized the disconnect between lesion burden and pain severity^22^.

This indicates the presence of uncovered mechanisms driving pain transmission. One leading hypothesis is that the neuro-immune signaling driven by local neuro-immune interactions underlies the increased nociceptive phenotypes and sustains chronic inflammation.

Despite the powerful insights gained from single cell approaches, these dissociative methods inherently disrupt tissue architecture, limiting our ability to study sensitive cell types or spatially integrated cell populations such as neurons and their projections. Even single-nuclei RNAseq fails to capture neurons whose cell bodies reside outside lesion boundaries, while low-resolution spatial transcriptomics (ex. Visium) lack the continuous capture area to uncover the small neurites found in lesion areas. The absence of high-resolution spatial information precludes direct inference of cellular neighborhoods, cell communities and cell-cell interactions within lesions. Thus, integrating single cell-resolution spatial information with single cell data is essential to resolve the multicellular architecture and signaling dynamics driving endometriosis.

Our previous work generated a comprehensive single cell atlas of endometriosis lesions, revealing coordinated immune phenotypes, increased angiogenesis, and similar lesion microenvironments^8^. Building on this foundation, we now extend the analysis to subcellular spatial resolution—placing cell types within their spatial context and defining spatial interaction networks that, critically, incorporate neurons as key components implicated in pain. This study: 1) creates a subcellular and continuous map of the cell type spatial architecture of human ovarian and peritoneal endometriosis lesions. 2) Identifies spatially defined clusters common across ovarian and peritoneal lesions explained by their specific fibroblast enrichment, their relative distance to epithelial glands and their neuronal enrichment. 3) Uncovers the spatially organized ensemble of sensory neurons and associated cell types — fibroblasts, immune cells, vascular components — likely engaged in bidirectional signaling that contributes to pain generation and sustained inflammation. Finally, 4) functionally validates the role of epithelial and *MME*+ fibroblasts in promoting sensory neurons, using in vitro 3D co-culture cellular models.

## Results

### Low-resolution spatial transcriptomics reveals spatial niche organization in ovarian and peritoneal lesions

We performed whole transcriptome spatial analysis using Visium v2 (10x Genomics) on three tissue types: endometrium biopsy (EMB), peritoneal superficial lesion, and ovarian lesion. The Visium assay is a low-resolution spatial assay that employs 55 µm spots to capture protein coding genes. These spots are spread out across the tissue with a 100 µm distance between the center of each spot, leaving substantial tissue space uncaptured (Fig. 1a). To identify epithelial glands – one of the key components of both EMB and lesions – we performed nuclei segmentation on H&E-counterstained tissues using QuPath and trained a learning model for gland detection. This model was integrated with spatial expression data for the epithelial marker *EPCAM* to define the spatial Epi cluster (Fig. 1a). In addition to the known presence of CD10+ (encoded by *MME*) fibroblasts, our previous endometriosis single-cell study led us to define different fibroblast subtypes in lesions. We specifically observed an increased osteoglycin (*OGN)* expressing fibroblast subpopulation in endometriosis tissues (EMB and lesions)^8^.

**Fig. 1:**
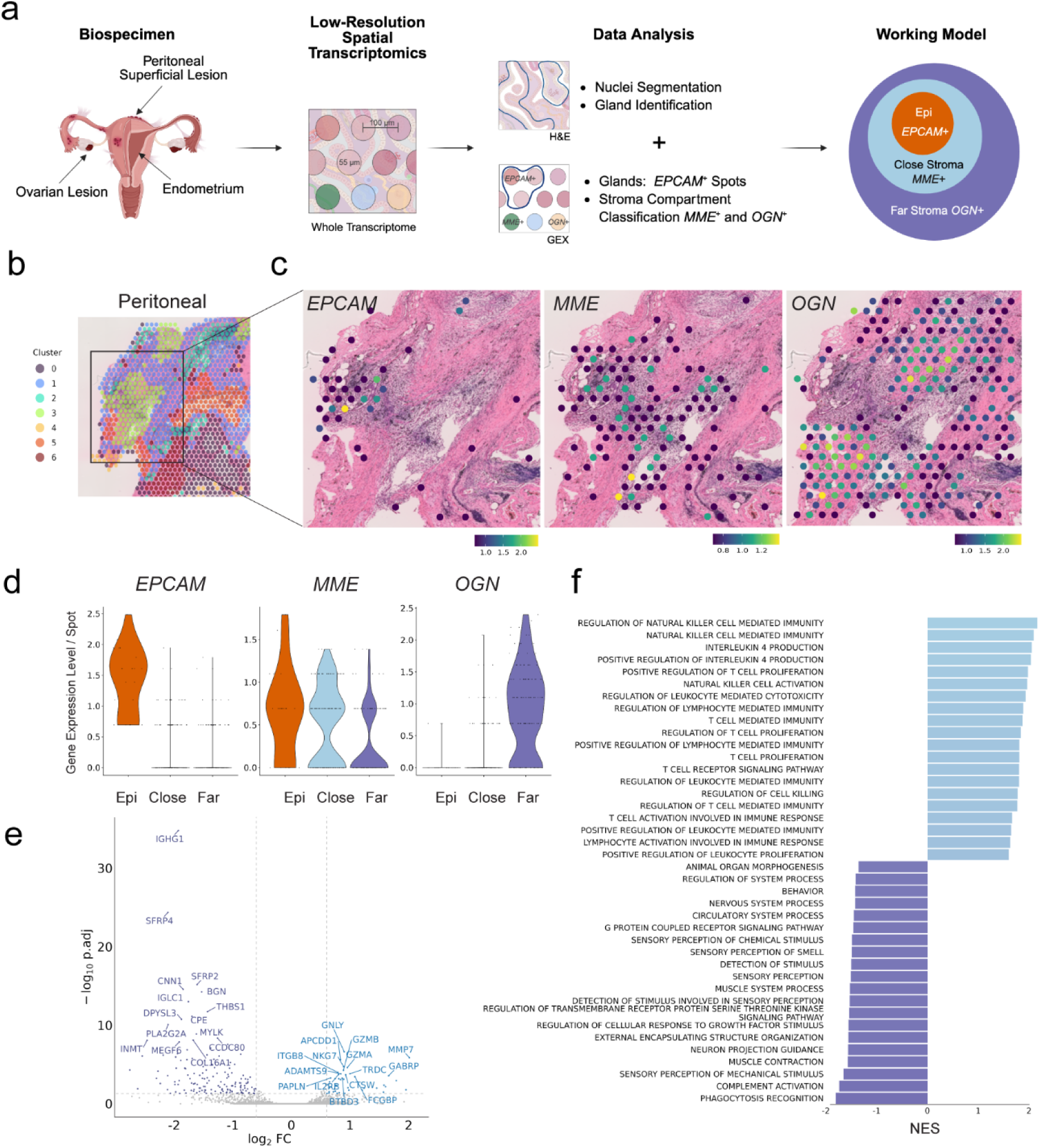
Low-resolution spatial transcriptomics of peritoneal endometriosis. **a**, Schematic of low-resolution spatial transcriptomics (Visium V2) workflow performed on peritoneal superficial lesion (n=1), ovarian lesion (n=1) and endometrium (n=1). Visium V2 is low-resolution due to 55 µM spots spread out across the tissue area, with 100 µM distance between the center of each spot, leaving substantial tissue space uncaptured. The H&E was used for nuclei segmentation and gland identification, the spatial transcriptomics data was used to identify *EPCAM*+, *MME*+, and *OGN*+ spots. The working model is as pictured with Close Stroma enriched in *MME*+ fibroblasts directly surrounding the epithelial glands and Far Stroma enriched in *OGN*+ fibroblasts outside of the Close Stroma cluster. **b**, Peritoneal lesion unsupervised clustering (sample P5). **c**, Zoom of identified gland area showing spots with detectable *EPCAM, MME*, and *OGN* expression. **d**, Violin plot of *EPCAM, MME*, and *OGN* expression levels in peritoneal Epithelial (Epi), Close Stroma (Close) and Far Stroma (Far) clusters. **e**, Volcano plot of the differentially expressed genes (DEGs) between the Close Stroma (blue, upregulated above zero) and Far Stroma (purple, downregulated below zero). The top 15 genes for each group are highlighted. The horizontal grey dotted line represents the significance threshold of p < 0.05 and the two vertical lines represent the Log2 fold change (FC) <-0.5 or > 0.5. **f**, Bar plot of top 20 significantly enriched GO: Biological Process pathways in the Close Stroma [blue, positive normalized enrichment score (NES)] compared to the Far Stroma (purple, negative NES).

Therefore, we investigated the spatial distribution of *MME* and *OGN* expression. Unsupervised clustering of the low-resolution spatial gene signatures failed to resolve the cellular complexity surrounding the glands, likely due to the heterogenous cell type composition within each spot, where we captured ∼25 cells per spot (Fig. 1b). We therefore manually annotated two stromal regions based on spatial proximity to epithelial glands and gene expression: the ‘Close Stroma’, an immediately adjacent region, marked by elevated *MME* expression, and the ‘Far Stroma’; and a more distal region enriched for *OGN* expression (Fig. 1c,d). This spatial stratification revealed a clear, layered pattern of *MME* and *OGN* expression in both ovarian and peritoneal lesions (Fig.1c,d) but not in EMB tissue (Supp. Fig.1a-f). These findings suggest the presence of lesion-specific stromal organization around epithelial structures.

To evaluate the molecular differences between stromal niches, we performed pseudo-bulking of the measured gene expression in Close and Far Stroma. We then conducted differential gene expression analysis (DEGs) within the peritoneal (Fig. 1e) and ovarian (Sup. Fig. 1c) lesions. In peritoneal lesion, the Far Stroma niche was enriched for genes associated with WNT signaling (such as *SFRP4*) and vascular processes signaling (*CNN1* and *THBS1)* while the Close Stroma displays elevated expression of immune cell, particularly NK cell, related genes, including *GZMA, GZMB*, *GNLY*, and *NKG7* (Fig. 1e,f). We observed enrichment of nervous system-related pathways in Far Stroma of peritoneal lesion (Fig. 1f). Though these findings were inconsistent in ovarian lesion (Sup. Fig. 1d), it became clear on further visual inspection of neuronal related that the platform lacked the sensitively to reliability detect fine neuronal processes such as small axons, as reported by Liu et al.^10^. To overcome this limitation and enhance spatial precision, we transitioned to high-resolution spatial transcriptomics using the Visium HD (10x Genomics) platform.

### High-resolution spatial transcriptomics allows for precise spatial clustering of lesion microenvironment

We performed high-resolution spatial transcriptomics using Visium HD on eight endometriosis lesions: four superficial peritoneal and four ovarian lesions (Sup. Table 1). We focused on these lesion types as they represent the two most prevalent clinical subtypes – peritoneal lesions are observed across all stages of disease while ovarian lesions affect ∼50% of patients undergoing treatment for fertility^23^. Visium HD enables high-resolution spatial mapping by capturing transcriptomic data across a continuous tissue area using 2 x 2 µm bins, leaving no tissue unsampled (Fig. 2a). We aggregated 4 x 4 squares of these 2 x 2 µm bins into 8 x 8 µm bins. This approach yielded a dataset comprising 2.62 million 8 x 8 µm bins, with an average of 327,000 bins per tissue section, providing an unprecedented subcellular spatial map of lesions (Sup. Table 2).

**Fig. 2:**
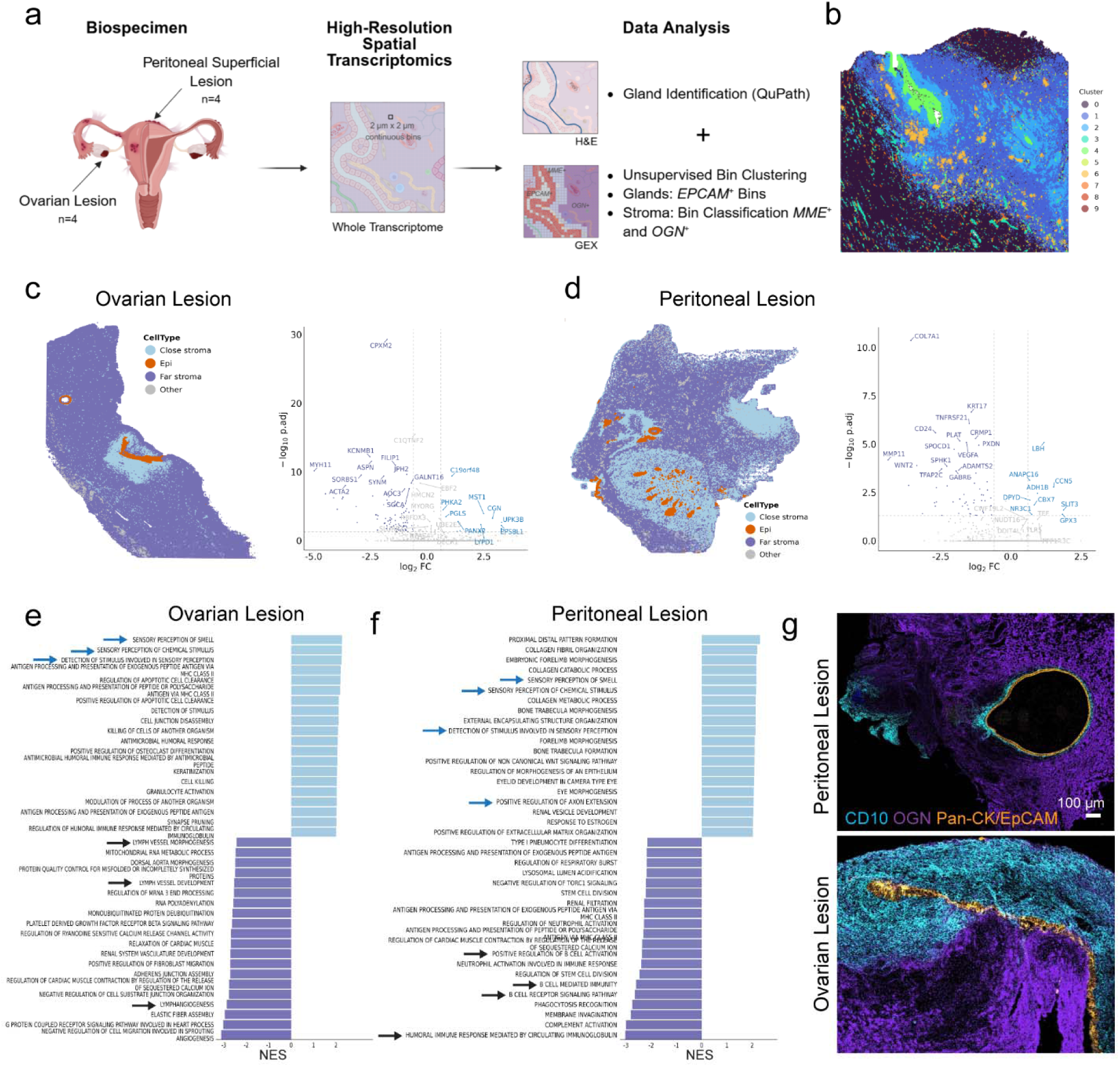
High-resolution spatial transcriptomics of ovarian and peritoneal endometriosis. **a**, Schematic of high-resolution spatial transcriptomics (Visium HD) workflow performed on peritoneal superficial (n=4) and ovarian (n=4) lesions. Visium HD is high resolution because it consists of 2 x 2 µm bins tiled continuously across the tissues area, leaving no tissue space uncaptured. The H&E was used for gland identification; the spatial transcriptomics data was used to identify *EPCAM*+, *MME*+ and *OGN*+ clusters. **b**, Unsupervised clustering of bins from **r**epresentative ovarian lesion--O2. **c,d**, (left) Representative image of ovarian lesion (sample O1a) (**c**) and peritoneal lesion (sample P1) (**d**) spatial clusters labeled as Epi, Close Stroma, Far Stroma or Other. (right) Volcano plot of top DEGs in the Close Stroma (blue, Log2FC > zero) compared to Far Stroma (purple, Log2FC < zero). The other six tissues can be found in Sup. Fig. 2. **e,f**, Bar plot of top 20 significantly enriched GO: Biological Process pathways in the Close Stroma [blue, positive normalized enrichment score (NES)] compared to the Far Stroma (purple, negative NES) for all ovarian lesions (**e**) and all peritoneal lesions (**f**). Arrows indicate pathways of interest. **g**, IMC protein validation of CD10 (encoded by *MME*) surrounding epithelial gland (Pan-cytokeratin/EpCAM) as Close Stroma and OGN farther away as Far Stroma. Scale bar is 100 µm. Ovarian lesion (sample O7) and peritoneal lesion (sample P8).

**Table 1:**
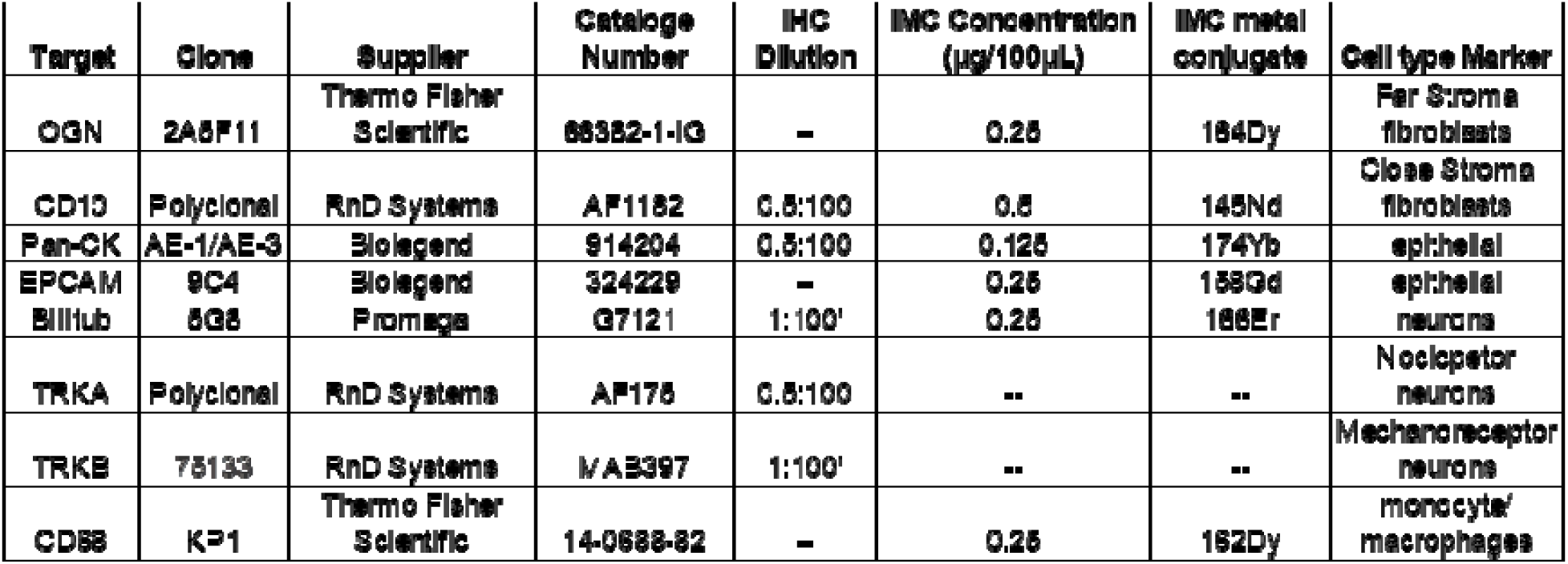
Antibody information.

Utilizing a similar method applied in our low-resolution spatial analysis, we identified epithelial glands using glandular feature detection on H&E images combined with spatial *EPCAM* expression across the bins. Unsupervised clustering of the transcriptomic profiles of 8 x 8 µm bins outlined distinct spatial niches (Fig. 2b). For each lesion, we labeled them based on mRNA expression of *MME* – for Close Stroma, *OGN* – for Far Stroma and *EPCAM* – for Epi (Fig. 2c,d). The Far Stroma included spatial clusters enriched in fibroblast-associated markers (*PDGFRA, LUM, SFRP4)* in addition to *OGN.* Clusters labeled ‘Other’ lacked fibroblast markers or represented low-quality bins. The corresponding protein expression relative to glands was confirmed by imaging mass cytometry (IMC) (Fig. 2g). The *MME/OGN* distribution pattern relative to the Epi niche was consistently found across all samples profiled (Sup. Fig. 2).

The Close and Far stroma gene expressions were subsequently pseudo-bulked for DEG and pathway analysis. Pathway analysis revealed distinct, cell type specific biological processes associated with each spatial cluster, consistent across lesion types. Pathways related to lymphatic vessels and B cells were significantly enriched in the Far Stroma in both lesion types (Fig. 2e,f, black arrows). Strikingly, sensory perception signaling pathways were enriched in the Close Stroma compartment across both ovarian and peritoneal lesions (Fig. 2e,f, blue arrows). This is in sharp contrast with findings from low resolution spatial results (Fig. 1f), underscoring the need for high resolution spatial profiling to resolve neuronal signatures accurately.

### Cellular proportions differ across stromal compartments and are consistent between peritoneal and ovarian lesions

To assess the full cellular complexity of the lesion microenvironment, we predicted the most likely cell type within each bin using marker genes from in-house and published single-cell datasets ^8–10^ (Fig. 3a,b, Materials and Methods section and Sup. Table 3,4).

**Fig. 3:**
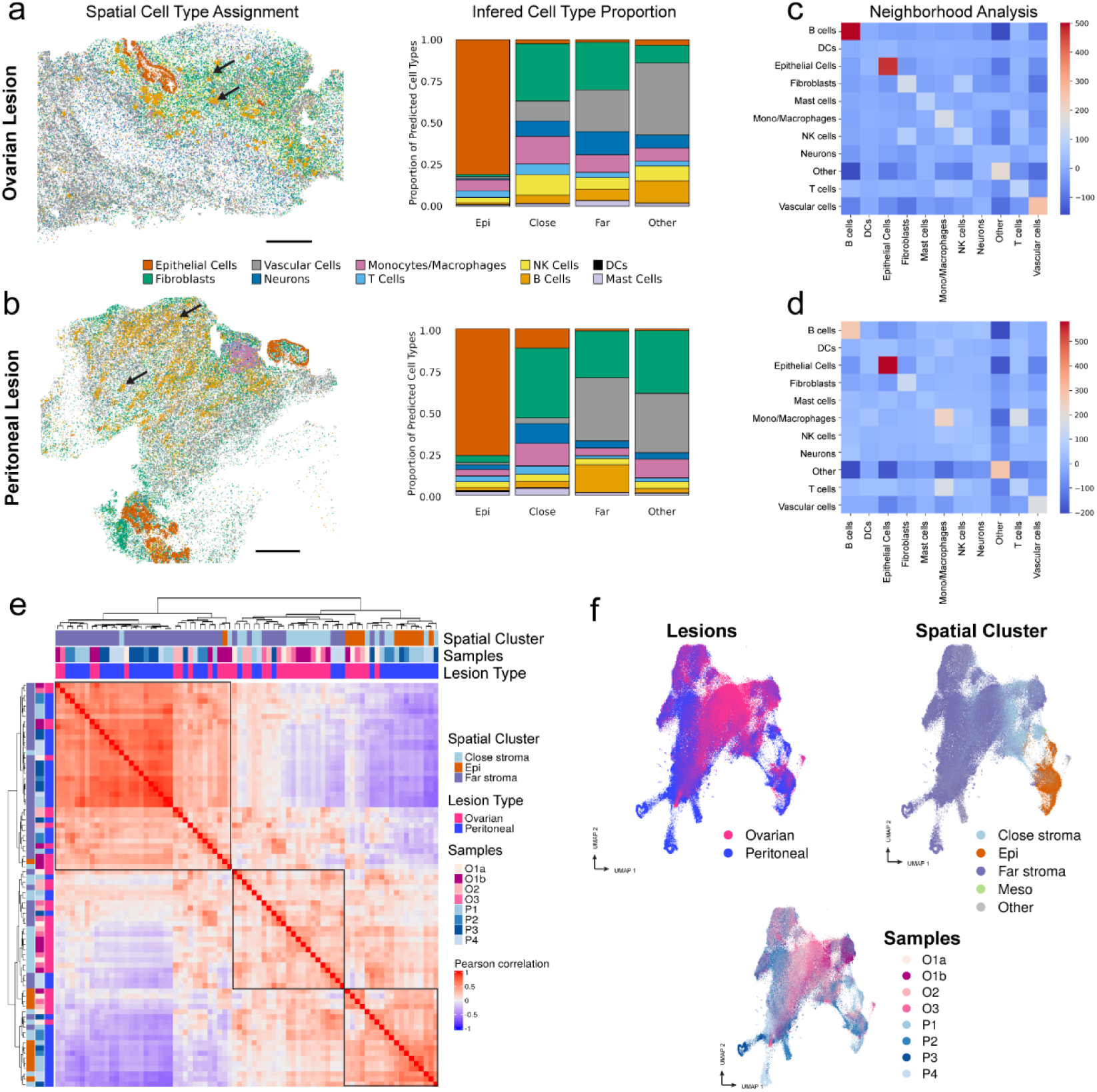
**Global cellular map of endometriosis lesions. a,b**, (left) Spatial plot of representative ovarian (sample O2) (**a**) and peritoneal lesion (sample P2) (**b**) showing bin-level cell type assignment for Epithelial Cells, Fibroblasts, Vascular Cells, Neurons, Monocytes/Macrophages, T Cells, NK Cells, B Cells, DCs and Mast Cells. Scale bar represents 1mm in **a** and **b**. (right) Bar plot represents the average proportion for each cell type in the Epi, Close Stroma, Far Stroma and Other spatially defined clusters for ovarian (**a**) and peritoneal lesions (**b**). Proportions for each tissue individually are in Sup. Fig. 3a. A linear mixed-effect model was used to compare cell type proportion differences across the stroma clusters in each lesion type; significant comparisons are shown in Sup. Fig. 3b. **c,d**, Neighborhood analysis heatmap of global cell types for lesions represented in panel (**a**) and (**b**). Per tissue analysis is found in Sup. Fig. 4. **e**, Heatmap of unsupervised clustering comparing spatial cell cluster, individual samples, and lesion type. Supervised clustering comparing peritoneal lesions and, separately, ovarian lesions is in Sup. Fig. 3c. Black boxes represent similar transcriptomic signatures in the spatially defined clusters. **f**, Uniform manifold approximation and projection (UMAP) of bins across all samples colored by lesion type (top left), spatial cell cluster (top right), and samples (bottom).

Shifts in predicted cell type proportions between our Epi, Close and Far Stromal spatial clusters showed similarities and differences between ovarian and peritoneal lesions (Sup. Fig. 3b). As a good control, epithelial cells were significantly enriched in Epi spatial clusters. Across both lesion types, vascular cells (which comprises endothelial and perivascular markers) are significantly enriched in the Far Stroma, and there is a trend towards increased monocytes and macrophages in the Close Stroma. Differences across lesion types were manifest in the significant enrichment of neurons within the Close Stroma of peritoneal lesions, while neurons were evenly distributed across Close and Far Stroma in ovarian lesions. NK cells are significantly enriched in the Close Stroma cluster of ovarian lesions (Fig. 3a,b, Sup. Fig. 3a,b).

B cells have been associated with endometriosis and autoimmune disorders involving B cells have also been linked to the disease^24^. B cell counts are increased in the peritoneal fluid of patients^25–28^, and we have observed tertiary-like structure (TLS) involving B cells in lesions^8^. However, their role remains unclear. Interestingly, we noted that B cells form aggregates throughout the lesion, and more abundantly in the Far Stroma in both lesion subtypes, across all tissues analyzed (Fig. 3a-d, Sup. Fig. 4).

To evaluate globally the similarities across lesion types, we integrated all tissues into one dataset and performed unsupervised hierarchal clustering on the pairwise similarities between all bins. The results indicate similar transcriptomic signatures across lesion types, mostly aligning with the spatial clusters initially defined independently within each sample by the stromal compartments (Fig. 3e, black boxes). UMAP projection of spatial-clustered bins shows discrete Close, Far and Epi clusters, with a little overlap, while lesion and sample UMAP plots indicate no sample bias (Fig. 3f). This further confirms our strategy to define spatial compartments and that the identified spatial features apply across both ovarian and peritoneal lesions.

### Neurons and immune cells, including macrophages, spatially organize within lesion

Neuro-immune signaling has been hypothesized to play an important role in the cell-to-cell interactions that occur in endometriosis lesions due to the association between increased inflammation and pain signaling^4, 17^. Thus, we evaluated the enrichment of immune cells in the immediate neuron periphery (within 16 µm distance) and compared it relative to all other bins (Fig. 4a). Results indicated increased NK-and T-cell enrichment around neurons for the Epi cluster, while B cells appear to be less present within the immediate neuron periphery in different spatial clusters. We also noted an enrichment in mast cells in proximity to neurons within both the Close and Far Stromal clusters and across peritoneal and ovarian lesions. This observation is consistent with prior studies reporting mast cells direct apposition to nerve fibers in lesions, where they are believed to contribute to nociceptive signaling through degranulation and release of pro-inflammatory mediators^29, 30^ (Fig. 4a). Interestingly, macrophages appear to be mostly enriched within the immediate neuron periphery in the Epi cluster. Crosstalk between macrophages and neurons is critical in endometriosis, as we and others have previously shown based on functional studies and in single cell analysis^8, 14, 17^. As such, we compared the distribution of distances between macrophages and neurons to that of distances between macrophages and non-neurons. Within spatial clusters, we showed a significant correlation between macrophage – neuron distance, normalized to all other cell types, indicating that macrophages and neurons position closer to each other, while not establishing immediate contact. This was particularly emphasized within the Epi and Far Stroma for ovarian and peritoneal lesions, respectively (Fig. 4b). These results were further confirmed through IHC as we saw CD68+ macrophages located close to βIIITub+ neurites (Fig. 4c).

**Fig. 4:**
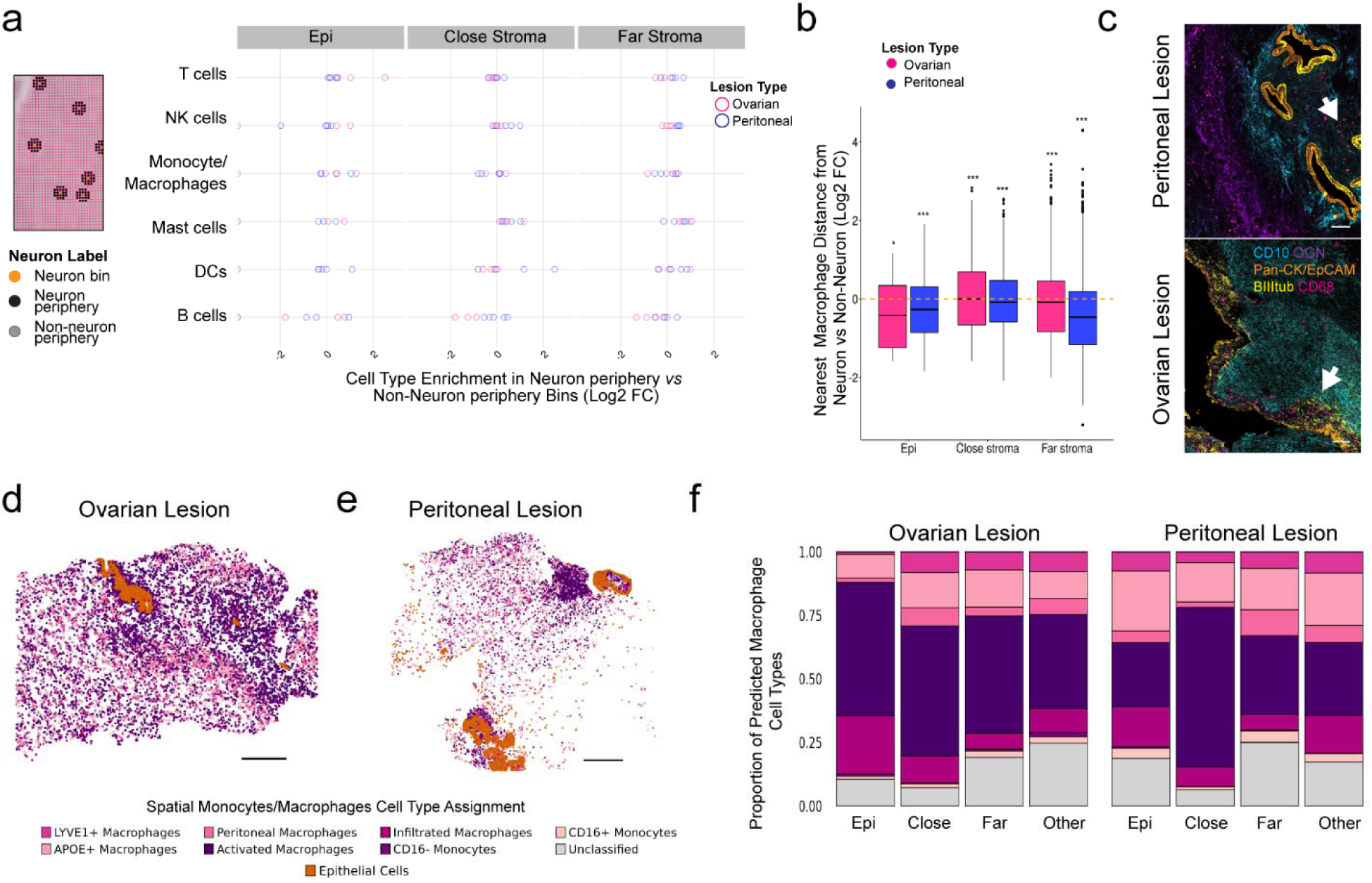
Spatial association of neurons to immune cells and macrophage subpopulations. **a**, Representation of bins annotated to neurons (orange) and two layers of bins (16 µm) surrounding a neuron bin labeled as neuron periphery (left). Per sample distribution of Log2 fold changes (FC) of immune cell types in neuron periphery compared to non-neuron periphery. Log2FC > 0 indicates immune cell enrichment in neuron periphery; Log2 FC < 0 indicates enrichment in non-neuron periphery bins. **b**, Box plot of distance calculation between a neuron and macrophage bin compared to a neuron and all other cell type bins. The box represents the span of the interquartile range (IQR), the middle line in the box is the median, the whiskers extend to the smallest and largest value of 1.5 x IQR, the points represent the outliers. Log2 FC is plotted, normalized to non-neuron bin distances to macrophages. Log2 FC < 0 (orange dotted line) have shorter distances between neurons and macrophages. Wilcoxon test adjusted p-values are plotted as: * p < 0.05, ** p < 0.01, *** p < 0.005. **c**, Antibody staining (IMC) indicates that neurons (βIII-tubulin+) are found located close to macrophages (CD68+) in proximity to glands, indicated by white arrows. CD10+ fibroblasts, OGN+ fibroblasts and Pan-cytokeratin/EpCAM+ epithelial cells were stained to visualize lesion area. Peritoneal lesion (sample P9) and ovarian lesion (sample O8). Scale bar in (**c**) represents 100µm. **d,e** Spatial plots of monocyte/macrophage subpopulations and epithelial bins. Representative image of an ovarian (sample O2) (**d**) and a peritoneal lesion (sample P2) (**e**). Dot size of monocyte/macrophage subpopulations and epithelial bins have been increased to visualize distribution patterns. **f**, Bar plot of average proportion of monocyte/macrophage subpopulations in ovarian and peritoneal lesions. Per sample proportions and linear mixed-effect model heatmap comparing average proportions in each lesion type are in Sup. Fig. 5.

Next, we examined the spatial localization of macrophage subtypes (Fig. 4d,e), which we previously identified in our endometriosis single cell atlas^8^. We observed a trend in increased presence of activated macrophages within the Epi and Close Stroma clusters for both lesions (Fig. 4f and Suppl. Fig. 5b). Ovarian lesions show a significant increase in LYVE1+ and peritoneal macrophages within the Close Stroma cluster (Fig. 4f and Sup. Fig. 5b). Altogether these results suggest a specific pattern of neurons and macrophages co-localizing in the endometriosis lesion microenvironment, where activated macrophages, expressing *SPP1, FN1* and *SPARC*, are specifically positioned near epithelial glands. Notably, *SPP1* expressing macrophages were reported to contribute to extracellular matrix (ECM) and tissue remodeling in endometriosis and other fibrotic diseases^31^.

While this spatial profiling does not provide mechanistic insights, it does confirm that macrophages and neurons are key partners in the endometriosis microenvironment. Our data further supports the concept of specific immune cell-neuron interactions that are not incidental but represent a conserved feature across endometriosis lesion types.

### *Identification of different sensory neurons subtypes: nociceptors sensory neurons spatially distribute close* to glands

We next examined the spatial distribution of neuronal subtypes, by subclassifying neuron bins (Fig. 3), expressing pan-neuronal markers (Fig. 5a), based on subtype-specific markers (Fig. 5b). Prior studies have reported increase of sensory neurons near endometriosis lesions^4, 5, 21, 32–35^, prompting us to evaluate specific sensory neurons subtypes: nociceptor (*TRPV1*, *NTRK1*) which respond to inflammation and transmit pain signals^14^, proprioceptor (*RUNX*, *NTRK3*) involved in body position sensing^36^, and mechanoreceptor (*PIEZO2*, *NTRK2*) which respond to mechanical stimuli^37^. We also included sympathetic neurons (*TH, SCL6A2*), down regulators of stress response^38^, and neuronal bins increased in immune-signaling related genes (*IL6R, TNFRSF1A),* labelled neuroimmune (Fig. 5a,b,c,f). 17% of the neuronal bins could not be subclassified, though they expressed pan-neuronal markers (Fig. 5a).

**Fig. 5:**
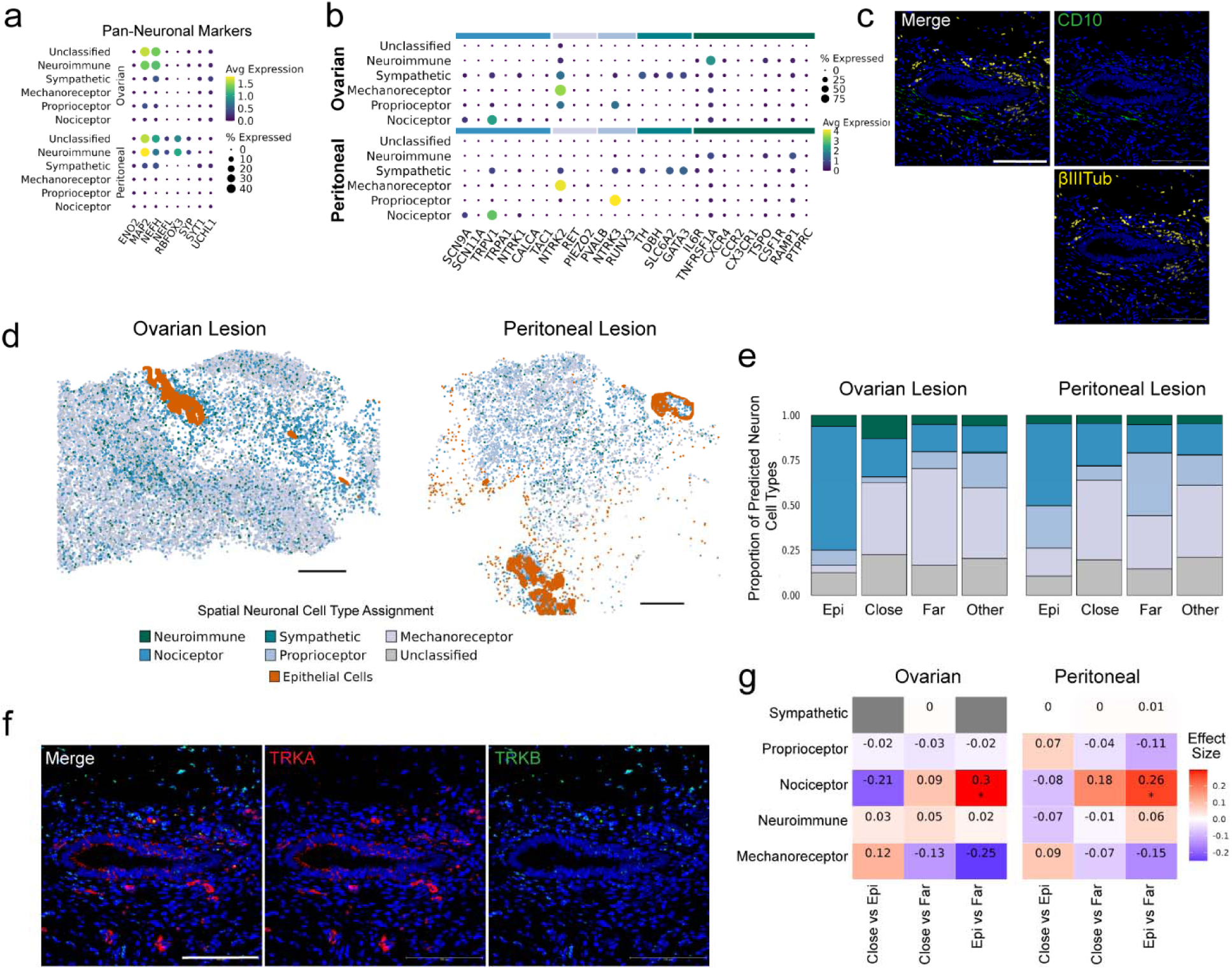
Spatial distribution of neuronal subtypes. **a**, Dot plot of pan-neuronal genes used to identify neuron bins. **b**, Dot plot of neuronal subtype gene marker expression in each neuronal subtype defined. **c**, IHC images of representative peritoneal lesion (sample P10) showing CD10+ fibroblasts and βIII-tubulin+ neurons surrounding epithelial gland. Scale bar represents 100 µm. **d**, Spatial plots of representative ovarian (sample O2; left) and peritoneal lesions (sample P2; right) showing the distribution of neuronal bins across the tissues, and dot size was increased to visualize distribution patterns. Epithelial cells are plotted to localize the lesion area. Scale bar in **d** represents 1mm. **e**, Bar plot of average neuronal subtypes in ovarian and peritoneal lesions across the spatial clusters. **f**, Protein detection of TRKA+ and TRKB+ neurons close to epithelial gland in a representative peritoneal lesion (sample P10). Scale bar represents 100 µm. **g**, Linear mixed-effect model heatmap of average neuronal subpopulation proportions in spatial clusters of ovarian and peritoneal lesions. The y-axis shows the cell types, and the x-axis shows the spatial cluster comparison. Number in each box is the effect size that correlates with the color scale, with red (positive effect) being increased in the first cluster compared and blue (negative effect) being higher in the second cluster compared. Stars represent significance: * p < 0.05.

Among neuronal subpopulations, we identified a significant enrichment of nociceptor neurons, marked by expression of pain receptors such as *SCN9A* and *TRPV1*, that localize within the Epi cluster in both ovarian and peritoneal lesions (Fig. 5d,e,g). On average, nociceptor neurons account for 18% of the neuronal bins across lesions. Ovarian lesions show a trend towards increased neuroimmune neurons – which express inflammatory chemokines such as *TNFRSF1A* and *RAMP1* – within the Close Stroma. Mechanoreceptor neurons, marked by *NTRK2* expression, show a trend toward enrichment in the Close and Far clusters across both lesion types (Fig. 5f,g). Notably, they represent the predominant neuronal subset accounting for an average of 40% of the neuronal bins. In contrast, sympathetic neurons are the least represented, comprising only ∼ 0.1% of the neuronal compartment. While we report average expression across lesions, individual tissues exhibit patient specific heterogeneity (Sup. Fig. 6).

Beyond nociceptive neurons, other sensory neurons such as proprioceptors and mechanoreceptors remain poorly characterized in endometriosis. Our data provides a novel, comprehensive overview of neuronal diversity within endometriosis lesions. Collectively, these results also reveal distinct spatial localization of neuronal subtypes, with sensory neurons – particularly nociceptors – enriched in the epithelial compartment.

### In vitro validation of neuronal dynamic interactions within endometriosis microenvironment

Given the preferential localization of the sensory neurons near the epithelial glands and adjacent to the Close Stroma compartment, we sought to functionally validate the dynamic interaction between epithelial, stroma cells – particularly *MME* expressing fibroblasts – and neurons. We generated 3D *in vitro* models recapitulating each of these components using: 1) peripheral sensory brain organoids (PSBOs) derived from a female iPSC line; 2) primary epithelial organoids derived from EMB and peritoneal and ovarian lesions; and 3) CD10+ fibroblasts derived from EMB and lesions, co-cultured in Matrigel domes (see Materials and Methods) (Fig. 6a). PSBOs were differentiated for 86 days and validated for the presence of sensory neuron markers TRKA, TRKB, TRKC (Fig. 6b). Co-cultures were maintained for 10 days and RNA was extracted for bulk RNA-sequencing to evaluate the effect of epithelial and stromal endometriosis cells on neuronal signaling pathways. Canonical makers for epithelial organoids (*KRT8, EPCAM*), fibroblasts (*PDGFRA, MME*) and mature neurons (*MAP2, RBFOX3, SYP*) were used to validate the experimental design and support that these co-cultures mimic the major cell types recapitulated in Epi/Close Stroma environments (Fig. 6c,e).

**Fig. 6:**
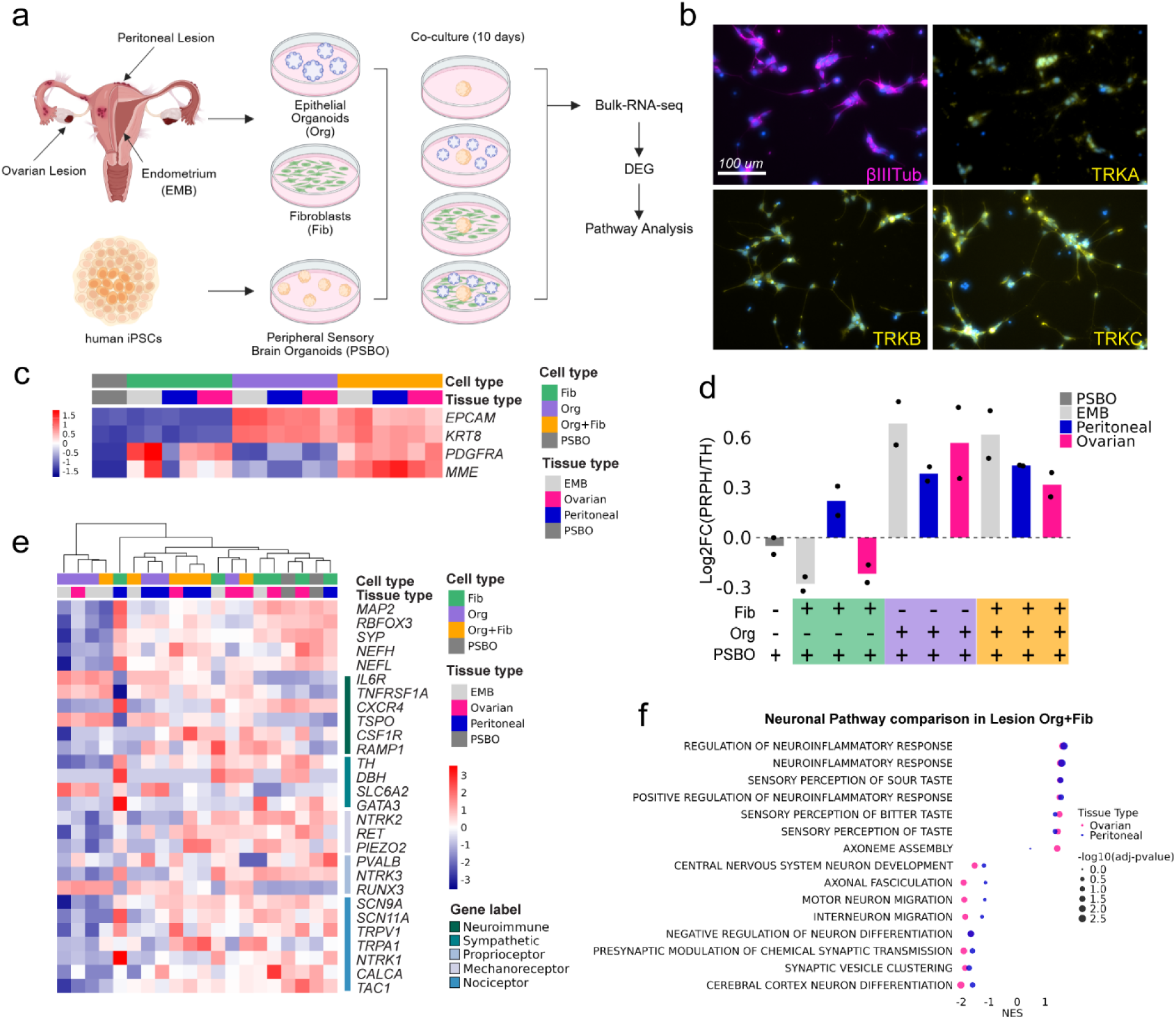
***In vitro* cell model of neuronal-epithelial-fibroblasts interactions in endometriosis. a**, Schematic of experimental design. Primary cell lines for epithelial organoids (org) and fibroblasts (fib) were derived from endometriosis patient endometrium, peritoneal or ovarian lesion tissues. Peripheral sensory brain organoids (PSBO) were derived from a female iPSC-line following the growth factor protocol outlined in the Materials and Methods section and Sup. Fig. 7. Cells were co-cultured for 10 days in different combinations for each tissue type: PSBO, PSBO+org, PSBO+fib, PSBO+org+fib. Bulk RNA-sequencing was performed on all the co-culture conditions (n=2). The resulting transcriptomics signatures were compared for DEGs and pathway analysis. **b**, IHC staining was preformed on dissociated PSBOs (differentiated to 28 days) confirming the successful differentiation of iPSC to neurons (βIII-tubulin), sensory nociceptor (TRKA+), mechanoreceptor (TRKB+), proprioceptor (TRKC+). Scale bar represents 100 µm. **c**, Heatmap showing the expression of epithelial markers (*EPCAM*, *KRT8*) and fibroblast markers (*PDGFRA*, *MME*) in analyzed co-culture conditions. **d**, Bar plot of the proprotion of PRPH+ sensory neurons over TH+ sympathetic neurons in all co-culture conditions, normalized to the PSBO alone condition. **e**, Heatmap of expression of genes used to identify *in vivo* spatial neuron subtypes, for the *in vitro* co-culture conditions. **f**, Dot plot of neuronal pathways significantly upregulated (NES > 0) or down regulated (NES < 0) in the ovarian and peritoneal co-culture PSBO+org+fib conditions.

We performed DEG analysis between the different conditions. PSBOs cultured alone differentiated towards *TH*+ sympathetic and *PRPH*+ sensory neurons with a distribution of 55: 45 percent. However, when the PSBOs were co-cultured with epithelial organoids (Org) or with both organoids and fibroblasts (Org + Fib), this proportion shifted to 40% for *TH*+ sympathetic and 60% *PRPH*+ sensory neurons (ratio in Fig. 6d). Fibroblasts alone were not able to drive this neuronal shift in PSBOs. These data suggest that the presence of epithelial organoids, endometrial and lesion-derived, drive neurons toward a sensory neuron fate.

We next compared the PSBO neuronal gene signatures to gene sets used to define the neuronal subpopulations in the spatial transcriptomics (Fig.6e). Neuroimmune genes *IL6R* and *TNFRSF1A* are both increased in the presence of organoids, while *RAMP1* and *CSF1R* seem to be driven by the presence of fibroblasts. We also observe the strong increased gene expression of inflammation-related genes, such as *NINJ1, IL1B, TNF, IL18* in Org+Fib co-cultures (Sup. Fig. 7). Mechanoreceptor genes, such as *RUNX3 and PVALB*, are upregulated in the Org+Fib conditions for all of the tissue-derived cells. Expression of nociceptor gene *TRPA1* is specifically upregulated in the peritoneal and ovarian lesion-derived organoid and fibroblast co-cultures compared to EMB co-cultures.

Finally, we performed pathway analysis between ovarian and peritoneal co-cultures for the PSBO/Org+Fib conditions. We extracted the pathways involved in neuronal signaling, showing that overall neuron differentiation was significantly increased across lesion type co-cultures. As well, genes associated with sensory neuron and neuroinflammation signaling pathways were increased in both lesion types, which further validates our spatial DEG analysis (Fig. 6f). We also noted that motor neuron and central nervous system signaling-related genes were down regulated in both lesion type-derived co-cultures. These data suggest that epithelial organoids in combination with MME+ fibroblasts push neurons toward a sensory fate and increase the neuroinflammation response signaling pathway alongside pain receptor expression, validating spatial transcriptomic findings.

### Ovarian follicle AMH levels are correlated with the distance from lesion epithelial glands

The link between infertility and endometriosis remains unclear, particularly how the presence of endometriosis lesions impacts the quality of ovarian follicles. We unexpectedly observed the presence of follicles close to the endometriosis lesion in one of the ovarian samples (sample O3). We identified follicles based on H&E staining and the expression of specific follicular markers (*ZP3*, *GDF9*,*OOSP2* and *FIGLA*)^39^(Fig.7a). Anti Mullerian Hormone (AMH), which is specifically expressed by small, growing follicles, is widely used as a marker for ovarian reserve and ovarian stimulation capacity^40^. Patients with higher AMH levels have been associated with better fertility outcomes^41^. All identified follicles were located in the Far Stroma (Fig.7b), with a variable range of *AMH* levels per follicle (Fig. 7c). We noted a significant positive correlation between a follicle’s *AMH* gene expression level and its distance from the epithelial gland (Fig.7d), suggesting that follicles may be impacted by the lesion. These data demonstrate the remarkable capacity of continuous high-resolution spatial transcriptomics to infer potential mechanisms associated with endometriosis infertility.

**Fig. 7:**
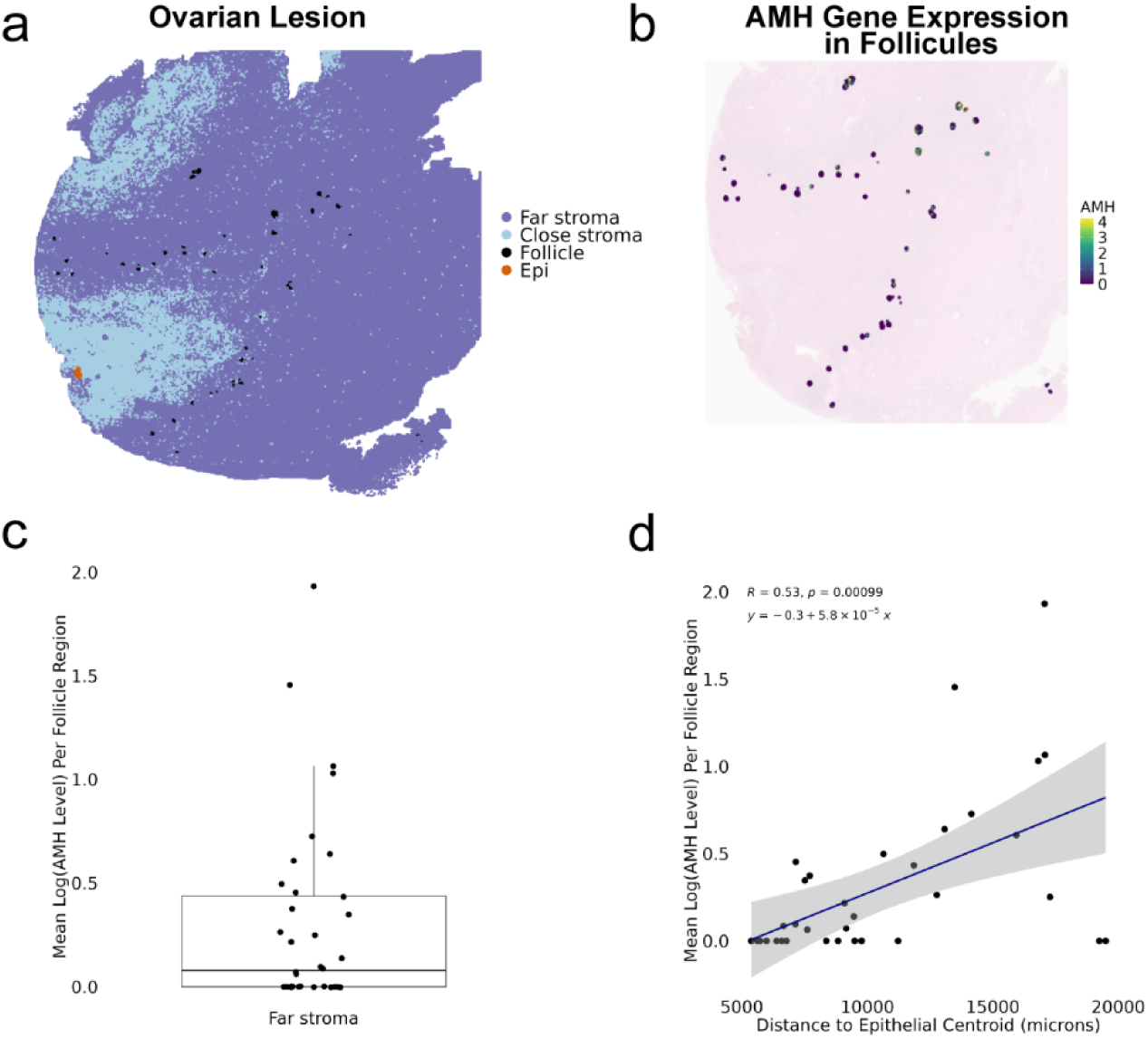
Ovarian follicle *AMH* level is correlated to distance from epithelial gland. **a**, Spatial plot of ovarian lesion (sample O3) with Epi, Close Stroma, Far Stroma and Follicles labeled. **b**, AMH expression in Follicle bins. **c**, Boxplot of average AMH levels per follicle in the Far Stroma. We did not identify follicles in the Close Stroma. **d**, Linear model comparing the follicle distance from the centroid of the epithelial gland to the average *AMH* level of the follicle. Blue line is the best fit linear regression model (y=-0.3+5.8×10^-5^x). Shaded grey is 95% confidence interval. The Pearson’s correlation was calculated to be R=0.53 and the p = 0.00099.

## Discussion

This study presents a high-resolution spatial transcriptomic profiling of human ovarian and peritoneal endometriosis lesions. Previously, we had established a comprehensive single cell atlas, based on dissociative approaches, of similar lesions. Yet this atlas lacked the spatial information needed to understand dynamic interactions and ecosystems driven by the spatial distributions of cells. In addition, due to technical limitations of dissociative approaches neurons, a known component of lesions and a key driver of endometriosis symptoms, were not detected. Now we have established a comprehensive spatial atlas of endometriosis lesions, at sub-cellular resolution, and provide an unbiased survey of the neuronal landscape within these structures. Although endometriosis is a highly heterogeneous condition, we employed spatially defined regions to identify shared pathways across two of the most prominent lesion types (Fig. 8). This approach emphasizes the importance of understanding the disease’s complexity and spatial patterning to identify common features and mechanisms among different lesion types.

**Fig. 8:**
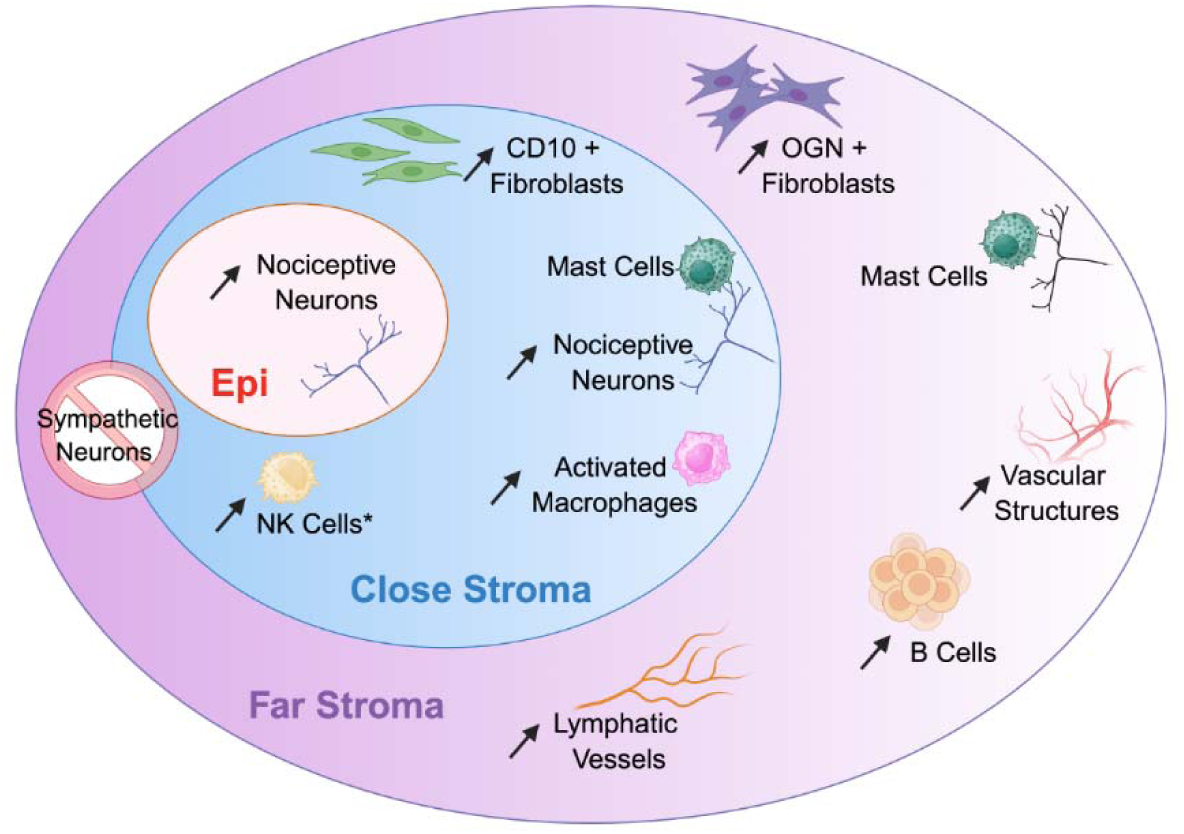
Representation model of common cellular features across lesions. Schematic of proposed model for spatial cluster cell type distributions in ovarian and peritoneal lesions highlighting common features across lesions. Epi Stroma cluster showed increased expression in Nociceptor neurons. The Close Stroma cluster showed increased expression of CD10+ fibroblasts, Nociceptor neurons, activated macrophages in both lesion types and NK cells in ovarian lesions only (indicated with *). The Far Stroma cluster showed increased vascular structures, B cells and lymphatic vessels in both lesion types. Globally, sympathetic neurons were sparse and there was a large increase in sensory neurons in the lesion microenvironment. Lastly, the data showed mast cells are enriched close to neurons in both the Close and Far Stroma clusters.

While spatial transcriptomics provides a powerful approach for inferring ecosystems that play a critical function in lesion pathobiology, it provides only a snapshot view of the dynamic cellular interactions underlying a complex microenvironment. Thus, we developed a 3D co-culture model, using patient derived primary lines, which recapitulates some of the critical cell types and their functional interactions to begin to understand dynamics between neuron-epithelial partners.

From our spatial transcriptomic analysis we defined shared spatial niches across lesion types, labelled Epi, Close and Far Stroma, based on mutual distance from glandular structures and shared expression signatures. Surprisingly, we found compelling similarities between spatial clusters of superficial peritoneal and ovarian lesions, two morphologically distinct lesions, with variable clinical stages and treatments. Our data support findings of a recent study on endometrioma spatial analysis, where spatial niches were also observed, with fibroblasts proximal to epithelial glands were implicated in wound healing whereas distal fibroblasts were defined as pro-inflammatory^10^. Liu and colleagues also determined that low-resolution spatial transcriptomics, while informative, does not provide sufficient clarity for comprehensive analysis of immune cells and neuronal organization. Indeed, we also demonstrated that low-resolution spatial transcriptomics did not reliably depict the positioning of immune and neuronal cells and only provided a discontinuous map of lesions.

Our spatial transcriptomic analysis showed the presence of B cell aggregates in the Far cluster consistently across all ovarian and peritoneal lesions. Previous reports have described an increased presence of peripheral and peritoneal fluid B cells in endometriosis patients, and showed that plasma cells produce autoantibodies against the endometrium and autoantibodies commonly found in autoimmune diseases^26, 42^. However, most of the B cell biology in female reproductive pathologies remain unknown^43^. Our work further suggests the contribution of B cell aggregates to the Far Stroma niche likely involved in promoting a chronic inflammatory environment. Parallels between endometriosis and autoimmune diseases, or immune-related disorders^24^, have been suggested in previous studies; however, our study further explores this perspective and identifies these aggregates as a common immunological mark across lesion subtypes, targeting of which could represent a promising therapeutic approach.

An unresolved question in the field is whether neurogenesis results from chronic inflammation or is driven by the presence of lesion-specific epithelial cells and fibroblasts in human tissues. With our data, we show an increase in nociceptor neurons expressing pain receptors close to epithelial glands across both peritoneal and ovarian lesions. Our *in vitro* 3D cellular model utilizing sensory brain organoids provides functional evidence of epithelial cells and fibroblasts in promoting sensory pain neurons. Further spatial analysis supports the increased presence of activated macrophages in areas where nociceptor neurons and epithelial glands colocalize. These data suggest a potential synergistic effect whereby epithelial cells influence neurons to express pain receptors. In turn, these neurons likely increase the production of inflammatory molecules (such as *NINJ1*), promoting the recruitment and activation of immune cells, particularly macrophages, and contributing to the sustained inflammatory microenvironment. Indeed, *NINJ1* was shown to mediate tissue remodeling and inflammation by regulating leukocyte entry, adhesion, activation, and migration^44^, and represents a potential candidate factor involved in endometriosis neuroimmune crosstalk. Further functional and mechanistic studies with complex 3D systems, recapitulating all these central cellular components will help elucidate the role of macrophages or mast cells in this process.

Endometriosis is an estrogen-dependent disease, and steroid hormone cyclic variation influences the immune landscape in reproductive female organs and related-diseases.

Technical and cost limitations of high-resolution spatial transcriptomics hinder the profiling of large clinical datasets. In this study we chose to focus on lesions from patients under hormonal contraception (HC) to minimize the impact of the menstrual cycle variation on our dataset. HC is also a standard treatment for endometriosis patients not seeking pregnancy. Interestingly, this study also suggests that HC may not be effective against the neurogenic and pain modulating processes driven by the presence of epithelial glands and close stroma. Additional studies have shown that estrogens play an important role in the mast and macrophage-neuron communication in endometriosis^30, 45^. Therefore, further research is necessary to evaluate the impact of HC on lesions in the context of neuroimmune signaling and to investigate how this may be altered in non-hormonally treated patients, and across menstrual cycle phases.

Finally, one limitation of spatial transcriptomics is that it encompasses only a single spatial modality. Future integration of spatial proteomics, ECM organization and other modalities such as lipidomics will provide additional insights, further enhancing our understanding of the complex microenvironment associated with endometriosis lesions. Spatial transcriptomics infers co-localization of cells but cannot prove direct cell-cell contact. Further *in vitro* cellular modeling is needed to truly address cell-cell contact and functional impact. Our initial spatial profiling focused on the two most common subtypes--superficial peritoneal and ovarian lesions.

However, considering the significant heterogeneity of endometriosis, it is essential to investigate all lesion subtypes, including deep infiltrating endometriosis and extra-pelvic lesions, in order to identify shared pathways that could contribute to the development of comprehensive treatment strategies for all lesion variants.

## Conclusion

By integrating whole transcriptome data with tissue morphology, spatial transcriptomics has revolutionized our understanding of diseases such as cancer and irritable bowel disease^46,47^. Here, we fill a critical gap in endometriosis research by revealing precise spatial cellular interactions within lesions and establishing parallels between two of the most common endometriosis lesions. We found that ovarian and peritoneal lesions share spatial features. Our findings highlight a prominent role for immune cells, uncover fibroblast compartments surrounding glands, and define the spatial distribution of neuronal and macrophage subsets. Understanding the cellular and molecular drivers of pain—typically among the earliest symptoms—may offer a path to preventing or mitigating the disease, particularly its most debilitating manifestations, and informing the development of novel therapeutic strategies.

## Materials and Methods

### Human endometriosis tissue collection

This study was approved by The Jackson Laboratory and University of Connecticut Health Center (UCHC) Institutional Review Board (IRB). The patients were enrolled either under The Jackson Laboratory IRB or the Endometriosis Research Innovation Support & Education (EndoRISE) IRB with written informed consent and consented to share recorded information.

Tissue specimens that were sourced from EndoRISE Connecticut (CT) Endometriosis Data & Biorepository. Individuals assigned female at birth, ages 18 and above, were consented and enrolled in the CT Endometriosis Data & Biorepository for longitudinal endometriosis research, agreeing to share biospecimens (tissues, whole blood, urine, peritoneal fluid) and phenotypic data. Tissues and phenotypic data (including quality of life, clinical and medical history data) were collected by endometriosis specialists at UCHC following the World Endometriosis Research Foundation (WERF) Endometriosis Phenome and Biobanking Harmonization Project (EPHect) guidelines and standard operating procedures. Specimens were immediately transported on ice to the biorepository at The Jackson Laboratory for processing and long-term storage (within 1 hour of collection). The EndoRISE Sample Access Review Committee (comprised of staff from UCHC and The Jackson Laboratory) reviewed the request for the data and biospecimen distribution for this project and approved the request unanimously. Patient information and use of specimens in this project can be found in Sup. Table 1.

Tissues were collected at UCHC and transported to The Jackson Laboratory on ice in MACS tissue storage solution (Miltenyi, Cat:130-100-008). Tissues were fixed in 10% Formalin in Phosphate Buffered Saline (PBS) for 24 hours at 4oC. Following fixation, tissues were washed three times in PBS and stored in 70% Ethanol at 4°C until sent to embedded in paraffin blocks. Formalin-fixed Paraffin embedded (FFPE) tissue blocks were sectioned at 5 µm and stained with Haematoxylin and Eosin (H&E) to visualize histology or Picrosirius Red (PS) to identify areas of dense collagen staining in lesions. For tissues where epithelial glands surrounded by fibroblasts were not obvious with the histological staining, we performed immunohistochemistry using anti-CD10 (R&D Systems, Cat:AF1182) to identify fibroblasts and anti-Pan-cytokeratin (Biolegend, Cat:914204) to identify epithelial glands as previously described^8^ (see below IMC, IHC and confocal imaging section).

### Tissue processing for spatial transcriptomics

Once a lesion area was identified with epithelial glands and CD10+ fibroblasts, a 6.5 x 6.5 mm region of interest(ROI) was selected for spatial transcriptomics. RNA quality control was performed on the selected FFPE blocks with the Qiagen RNAeasy FFPE kit (Cat: 72504) followed by Agilent tapestation analysis of RNA. Only FFPE blocks with a DV200 above 30% were used for spatial transcriptomics.

FFPE blocks that passed all quality control metrics were sectioned at 5 µm and put on glass slides. The sectioned tissues were either processed following the 10X Visium V2 protocol or Visium HD V1 protocol by the Single Cell Biology Core at The Jackson Laboratory. Visium V2 was run on three samples, eutopic endometrium (EMB), peritoneal lesion and ovarian lesion, from two patients. Visium HD was run on eight samples, four peritoneal lesions and four ovarian lesions, from seven patients. Sup table 2 includes tissue information and spatial metrics.

### Spatial transcriptomics pre-processing and cluster identification

The Illumina base call libraries were converted to FASTQ files. The 10x Genomics Space Ranger pipeline (v3.0.0) was used to align the sequencing reads to the human reference genome GRCh38-2020-A. We keep the spots that express more than 100 genes and have less than 10% of mitochondrial gene counts. QuPath was used to segment the nuclei in the H&E image. The transcriptional data was used to manually label the spots containing EPCAM+ epithelial glands, MME+ close fibroblasts and OGN+ far fibroblasts.

For tissues processed with Visium HD V1, the resulting count matrices were processed in R using the Seurat package using the 8 x 8 µm binning. For each tissue individually, 8 µm bins were removed outside of the refined tissue mask, and 8 µm bins were removed with a UMI count less than the threshold that separates the foreground and the background of the refined tissue mask. The H&E was used to identify the epithelial glands, but unsupervised clustering was used to identify the fibroblast compartments. The clusters closest to the glands and highest in *MME* expression were labeled as ‘Close Stroma’. The clusters outside of the *MME*+ areas and higher in *OGN* were labeled as ‘Far Stroma’. The left-over clusters that did not fit this definition were labeled as ‘Other’.

### Visium HD sample integration and clustering

Due to large sample size (Sup. Table. 2) sample integration was performed on a representative set of cells from each sample. This sampling approach is based on a geometric sketching method, and it ensures that rare populations are also included in the subset^48^. The integration of eight samples was done with Seurat using the Harmony method on Principal Component space. Then, the integration was expanded to the full dataset.

An aggregated pseudo-bulk-based workflow was used, where each cell is aggregated sample of one cluster in each sample. Then clustering was done using the complete method and Euclidean distance (Fig. 3e). After cell aggregation, cell type specific differential expression was performed using DESeq2 method and Wilcoxon rank sum test. Pathway analysis for the biological process gene ontology terms (GO:BP) was run.

### Visium HD Cell type assignment and distance analysis

To predict the major cell type within each bin, Seurat’s AddModuleScore function was used with known markers for the following cell types: Epithelial Cells, Fibroblast Cells, Vascular Cells, Neurons, Macrophages, NK cells, T Cells, B Cells, DCs and Mast Cells (Sup. Table 3). The resulting scores were scaled and filtered for a z-score above 1 to be labeled as a cell type. The spatial location of all the cell types was plotted using SpatialDimPlot and increased the point size to visualize the patterns. The proportion of each cell type per sample in each defined fibroblast compartment (Epi, Close or Far) was calculated by taking the number of bins for that cell type in the fibroblast compartment divided by the total number of bins in that fibroblast compartment and graphed it in a proportion plot. To assess which cell type proportions were significantly different between the stroma compartments across lesion types, post-hoc analysis was performed of linear mixed-effect models fit independently for each cell type and lesion type, with compartment as the fixed effect and sample as the random effect (Sup. Fig. 3b)

To identify subpopulations of neurons, all the bins labeled as neurons from the previous global cell typing were subsetted and run through AddModuleScore with another set of genes specific to the subpopulations (Sup. Table 3). Again, the scores were scaled but a z-score above 0.01 was used to identify the subpopulations of neurons as these genes are often low in abundance. The same process was performed to identify macrophage subpopulations using the gene markers from a published single cell dataset of endometriosis tissue^8^ (Sup. Table 3). The same linear mixed-effect model was used to calculate the differences between the subtypes of neurons and macrophages in the fibroblast compartments (Sup. Fig. 5b & Fig. 5g).

### Enrichment analysis between neurons and immune cells

To check whether immune cells are enriched around neuron bins, the proportion of each immune cell type in the bins directly surrounding the neuron bin, defined by two layers of 8 x 8 µm bins (neuron periphery in Fig. 4a), was calculated. The proportion of each immune cell type around all the non-neuron bins was calculated (non-neuron periphery in Fig. 4a). These values were normalized to show their fold change.

To evaluate the enrichment of macrophages in the vicinity of neurons within each compartment (Epi, Close Stroma, and Far Stroma), the distance was calculated between each neuron bin in that compartment and its closest macrophage, noting that the neuron and macrophage bins may not fall in the same compartment. Similarly, the distance between each non-neuron, non-macrophage bin in each compartment was calculated relative to its closest macrophage bin. The adjusted p-value was graphed from the result of a Wilcoxon test corrected using Benjamini-Hochberg procedure. Within each sample, all distances were log normalized to the median of the non-neuron/non-macrophage to macrophage distances within that sample (Fig. 4b). As such, a log normalized distance less than zero indicates that a macrophage is closer to a neuron than the median distance between a macrophage and a non-neuron, non-macrophage.

### IMC, IHC and confocal imaging

We used previously published Imaging Mass Cytometry (IMC) data reanalyzed for markers specific to this project (Fig.2 & 4)^8^. Immunohistochemistry (IHC) was adapted from previously described protocol^8^ for the localization of TRKA and TRKB neurons (Fig. 5). Briefly, FFPE tissue sections were melted for 10 minutes at 55°C, deparaffinized in 2 washes of Histoclear (National Diagnostics, Cat: HS-200). The tissues were rehydrated in a gradient of ethanol to water washes. Antigen retrieval was performed in EDTA pH 9 (Vector Labs, Cat: H-3301-250) for 15 min at 95°Cin a pressure humidity chamber (BioSB TintoRetriever). Sections were permeabilized in 0.1% triton for 10 minutes, then blocked in 10% donkey serum and 3% BSA solution for 1 hour at room temperature. Primary antibodies were incubated in 3% BSA in PBS overnight at 4°C. Table 1 includes primary antibody information. Unbound primary antibody was washed away, then sections were incubated in secondary antibodies for 2 hours at room temperature. Secondary antibodies were used at 1:1000 dilution, Donkey Anti-Goat IgG-Alexa fluor 647 (Invitrogen, Cat: A32849) and Donkey anti-Mouse IgG-Alexa Fluor 555 (Jackson Immunological, Cat: AB_3095480). A Dapi counter stain was used. Slides were coverslipped with Flouramount-G (Life Tech, Cat:00-4958-02).

IHC images were acquired using a Leica TCS SP8 laser scanning confocal microscope (Leica Microsystems, Wetzlar, Germany) equipped with a DM6000 upright microscope stand. The system was controlled by Leica Application Suite X (LAS X) software (version 3.5.7.23225). Images were obtained using the HC PL APO CS2 40x/1.30 Oil objective (Cat #: 11506358).

Leica Microsystems’ Type F immersion oil was used (Cat #: 11513859). Excitation was provided by a combination of a 405 nm diode laser and a White Light Laser (WLL). Laser power was controlled via the software and was typically set between 1-10% to minimize photobleaching.

Fluorescence emission was detected by a combination of photomultiplier tubes (PMTs) and high-sensitivity hybrid detectors (HyDs). The spectral detection bands were tuned to optimally capture the emission of each fluorophore while minimizing crosstalk. The spectral detection bands were tuned as follows:

- DAPI: Excitation at 405 nm, emission collected at 415-475 nm.
- Alexa Fluor 488: Excitation at 499 nm, emission collected at 509-572 nm.
- Alexa Fluor 568: Excitation at 577 nm, emission collected at 587-648 nm.
- Alexa Fluor 647: Excitation at 652 nm, emission collected at 664-778 nm.

Images were acquired as 8-bit files with a frame size of 512 x 512 pixels. A conventional galvanometric scanner was used. For imaging, a scan speed of 400 Hz was used with a line averaging of 1. The pinhole was set to 1 Airy unit (AU) to ensure confocality. For Z-stacks, images were acquired with an average step size of approximately 2.0 µm, covering a total axial distance of approximately 8.0 µm. Sequential scanning was employed for multi-channel imaging to prevent bleed-through between channels. The scan sequence was set to “between frames”. Raw image files (.lif) were processed and analyzed using LAS X software and Fiji/ImageJ. The captured signals were max projected.

### AMH-follicle distance analysis

We received one lesion that came from an ovarian cortex and contained follicles near an endometriosis lesion (Fig.7). The Seurat function SpatialFeaturePlot was used to plot the AMH level for the follicle cluster on the tissue. K-nearest-neighbor was used to spatially localize the follicles into the closest stroma cluster. To analyze the quality of those follicles, the centroid of the epithelial gland and each follicle were calculated. DBscan was used to detect the bins for each follicle resulting in 35 follicles and the average AMH for each follicle was calculated. The Euclidean distance from the epithelial gland centroid to the follicle centroid was compared to the mean AMH level for each follicle using a linear regression model.

### Endometrial epithelial organoid and fibroblast culture

Tissues were dissociated into single cell suspension as previously described^8^. The single cell suspension was resuspended in cold Matrigel (Corning, Cat:356231) in 50 µL domes in a 24-well plate. Epithelial organoid media, previously described^49^, was added to cover the domes. For the first passage, to separate the epithelial organoids from the fibroblasts, the fibroblasts were allowed to grow to the bottom of the dish, then the Matrigel dome was carefully removed to leave the fibroblasts in place. The fibroblasts were washed with DPBS and fibroblast media was added. Fibroblast media contained DMEM/F12 (Gibco, Cat:11320082) supplemented with 10% dextran-coated charcoal stripped FBS (Sigma, Cat:C6241, Gibco, Cat:A5256801), 1% penicillin-streptomycin, 2 mM L-glutamine, 1 nM β-Estrodiol (Sigma, Cat:E4389), 2 ug/mL insulin (Sigma, Cat:91077C). Epithelial organoids were passaged every 7 days according to previously described protocol^49^. Fibroblasts were passaged 1:4 every 7-10 days once 80% confluent with a five-minute digestion of TrypLE (Gibco, Cat:1260510) to lift the cells from the plate.

### Peripheral Sensory Brain Organoid culture

Human induced pluripotent stem cells (hiPSCs) were cultured and induced to brain organoids adapted from the following protocols^50, 51^. All material and reagents can be found in the protocols.io (dx.doi.org/10.17504/protocols.io.kxygxw1ddv8j/v1). The hiPSCs were derived from a 24 year old healthy female of African ancestry at the Jackson Laboratory (GeDiX697.3J). This hiPSC line was fully validated for trilineage differentiation potential and whole genome sequencing. Briefly, to induce neuronal progenitor cells in organoid formation, the dual-SMAD method was adapted as previously described^52^ on a shaking platform at 95 rpm inside a 37oC incubator. Briefly, cells were cultured in 100% KSR medium supplemented with 10μM SB431542 and 1μM LDN193189 for days 4-8 post seeding. To promote the differentiation to sensory neurons a protocol was adapted^53^, 2.5 μM SU5402, 1 μM DAPT and 1.5 μM CHIR99021 was added on days 5-13 of differentiation. The cells were transitioned from 100% KSR media to 100% Neural Induction Media (NIM) over a 7 day gradient from day 8-14. Once in 100% NIM on day 14, the supplements were switched to 10 ng/mL FGF2 and 10 ng/mL EGF2 for 6 days, washing every other day. The cells were finally put into Terminal Differentiation (TD) media supplemented with 20 ng/mL BDNF, 20 ng/mL GDNF, 25 ng/mL NGF, 10 ng/mL NT-3, 200 µM Ascorbic Acid and 0.5 mM dbcAMP on day 20 post feeding. The cells were washed every 3 days with fresh media and supplements (Sup. Fig. 8).

To verify the neural induction to peripheral sensory organoids, the brain organoids were dissociated at 28 days in accutase for 20min at 37oC. Gentle pipetting was done to manually dissociate the organoids. The accutase was stopped by adding 3x the volume TD media. Cells were spun down 300 g for 5 minutes at room temperature. The cells were counted and viability checked, then 200K cells were plated per well in a 24-well dish, pre-coated with Matrigel, with TD media supplemented with growth factors. After 24 hours, the neurites grew out and the cells were fixed in 4% PFA for 2 hours at room temperature. The fixed cells were then stained for anti-β-III-tubulin (Promega, Cat: G7121), TRKA (R&D Systems, Cat:AF175), TRKB (R&D Systems, Cat:MAB397), and TRKC (R&D Systems, Cat:AF373) to detect different sensory neuron populations. The immune fluorescence protocol was as follows: cells were washed 3 times with PBS, permeabilized with 0.25% Triton-X100 in PBS for 15 minutes, washed 3 time with PBS, blocked with 10% donkey serum, 1% BSA in PBS for 1 hour room temperature, and incubated in primary antibodies at 1:100 dilution in 0.1% Triton-X100, 1% BSA in PBS over night at 4oC. On the second day, cells were washed 3 times for 5 minutes in PBS, incubated in secondary antibodies 1:1000 for 2 hours room temperature, washed 3 times for 5 minutes with PBS, incubated with DAPI for 5 minutes, washed 3 times for 5 minutes with PBS. Cells were imaged on Nikon Eclipse inverted fluorescent microscope at 20x magnification. Images were optimized for brightness and contrast across the entire field of view with FIJI.

### Co-culture and bulk RNA-seq of brain organoids, epithelial organoids and fibroblasts

To assess the molecular changes that occur when endometriosis epithelial cells and fibroblasts are in contact with sensory neurons, co-culture experiments were performed. Eutopic endometrial, peritoneal lesion and ovarian lesion epithelial organoids and fibroblasts were cultured in different combinations (denoted in Fig. 7a) with 84 day sensory peripheral brain organoids. After 10 days of co-culturing, the Matrigel domes were melted on ice for 30 minutes, the cells were spun down 520 g for 5 minutes at 4oC, cells were washed with PBS once and spun down again. The cells went right into RNA extraction following the Qiagen RNeasy Mini Kit protocol (Cat: 74106). RNA quality was assessed with Aglient Tapestation and only samples with a RIN above 6 were used for bulk RNA sequencing. Samples were prepped for whole genome library prep. Then sequenced on Novaseq X Plus 10B with 300 cycles with an average of 2.9 million reads per sample.

### Analysis of bulk RNA-seq

Reads were aligned with the human genome GRCh38-2020-A, filtered with nf-core/rnaseq (v.3.19) utilizing the STAR aligner^54^ and counts were normalized to counts per million. DeSeq2 was used to subtract the brain organoid alone conditions from all the rest of the conditions to get DEGs. The VST scaled matrix was used to assess gene expression differences between groups. Then the different co-culture conditions were compared using FGSEA pathway analysis. The neurona-related pathways were selected to compare between groups.

### Figure creation

All figures were put together using Adobe Illustrator. All schematics were made using Biorender.

## Data Availability

The data will be deposited on the public data repository such as Gene Expression Omnibus (GEO) upon publication. All other data for this study will be provided by the corresponding author upon reasonable request.

## Code Availability

All code developed and used for this study will be available on a GitHub page upon publication.

## Acknowledgements

The authors would first like to thank all of the study participants that generously donated their tissues to help us better understand the biology of endometriosis. Without their participation, none of this would be possible. We heartfully thank the EndoRISE Biorepository for enabling part of this research through access to biospecimen, and in particular Kayceety Mullaj, Makayla Murphy, and Jasmina Kuljancic for their support. Next, we would like to thank all of The Jackson Laboratory core facilities that supported this work and in particular: Genome Technologies, Histology, Microscopy and Cyberinfrastructure High performance computing resources. Of particular importance, we gratefully acknowledge the advanced expertise of the Single Cell Biology Lab and their continuous support. These shared services are supported in part by the JAX Cancer Center (P30 CA034196). We thank the OB/Gyn department and UConn Health Surgery Center personnel for their assistance in consenting patients and collecting specimens. A special thank you to the Robson and Skarnes Labs at the Jackson Laboratory for the use of iPSC cells, especially J. Alcoforado Diniz and C. Loring for their guidance, and A. Melo Carrilo for his help with the bulk-RNA-seq analysis. This study was supported by The Mayday Fund. C.M.H. is supported by Mayday Fund. E.T.C, J.K, and D.E.L are supported by the State of Connecticut, through HB6672 which funds the CT Endometriosis Tissue and Data Biorepository, which is part of the EndoRISE program. The iPSC line used in this study was derived with support from The Jackson Laboratory, Director’s Innovation Fund, award number DIF-FY22-Skarnes-iPSCDiversity.

## Contributions

E.T.C conceived and designed the study. C.M.H. and E.T.C performed and supervised experiments. D.E.L and J.K. collected consent from participants and clinical samples. C.M.H., J.L., J.K, E.T.C processed and biobanked samples for downstream experiments. J.K. provided EndoRISE patient information. M.S. and C.M.H. performed IHC experiments. J.L. imaged IHC.

C.M.H. conceived and performed *in vitro* experiments and bulk-RNA seq. C.M.H and W.F.F performed bulk-RNA-seq analysis. P.R. provided knowledge and resources for iPSC brain organoid cultures. C.M.H, E.A., and B.S.W. performed data analysis. B.S.W, W.F.F, and E.T.C., provided crucial data interpretation. C.M.H and E.T.C. wrote the manuscript and generated figures and schematics. All authors reviewed and edited the manuscript.

## Ethics declaration

The authors declare no competing interests.

