## Supplemental Figures for "High-Resolution Spatial Transcriptomics Reveals Fibroblast and Neuroimmune Microenvironments in Endometriosis Lesions"

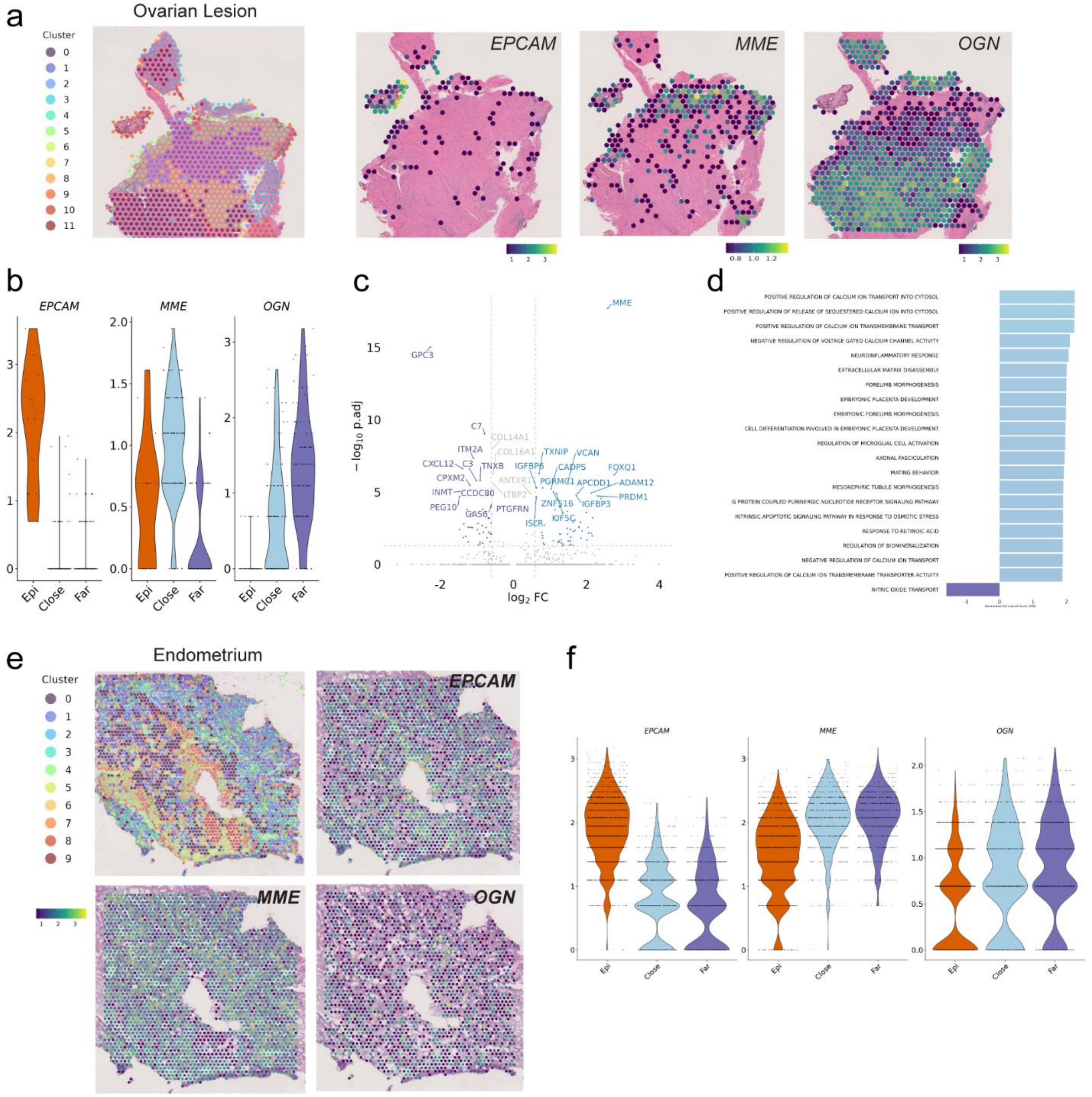


**Sup. Fig. 1: Low-resolution spatial transcriptomics of ovarian lesion and endometrium.** **a**, Ovarian lesion -- O4 unsupervised clustering and identified gland area showing spots with detectable *EPCAM, MME,* and *OGN* expression. **b**, Violin plot of *EPCAM, MME,* and *OGN* expression levels in ovarian Epithelial (Epi), Close Stroma (Close) and Far Stroma (Far) clusters. **c**, Volcano plot of the differentially expressed genes (DEGs) between the Close Stroma (blue, upregulated above zero) and Far Stroma (purple, downregulated below zero) within the ovarian lesion. The top 15 genes for each group are highlighted. The horizontal grey dotted line represents the significance threshold of $p < 0.05$ and the two vertical lines represent the Log2 fold change (FC) < -0.5 or > 0.5. **d**, Bar plot of top 20 significantly enriched GO: Biological Process pathways in the Close Stroma [blue, positive normalized enrichment score (NES)] compared to the Far Stroma (purple, negative NES). **e**, Endometrium, Emb1, unsupervised clustering and spots with detectable *EPCAM, MME,* and *OGN* expression. **f**, Violin plot of *EPCAM, MME,* and *OGN* expression levels in endometrium Epi, Close and Far clusters.

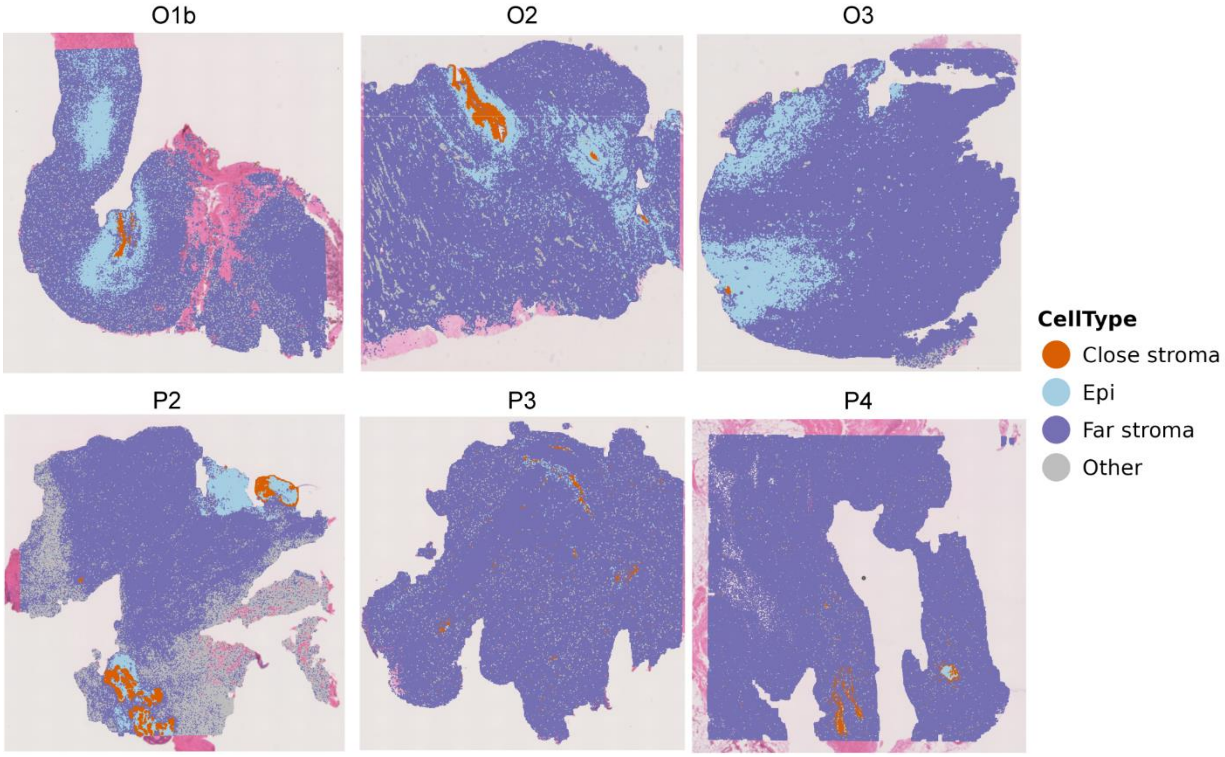


**Sup. Fig. 2: Close and Far Stroma clusters for each high-resolution ovarian and peritoneal lesion.**
Spatial plots of ovarian lesion (top) and peritoneal lesion (bottom) clusters labeled as Epi, Close Stroma, Far Stroma or Other.

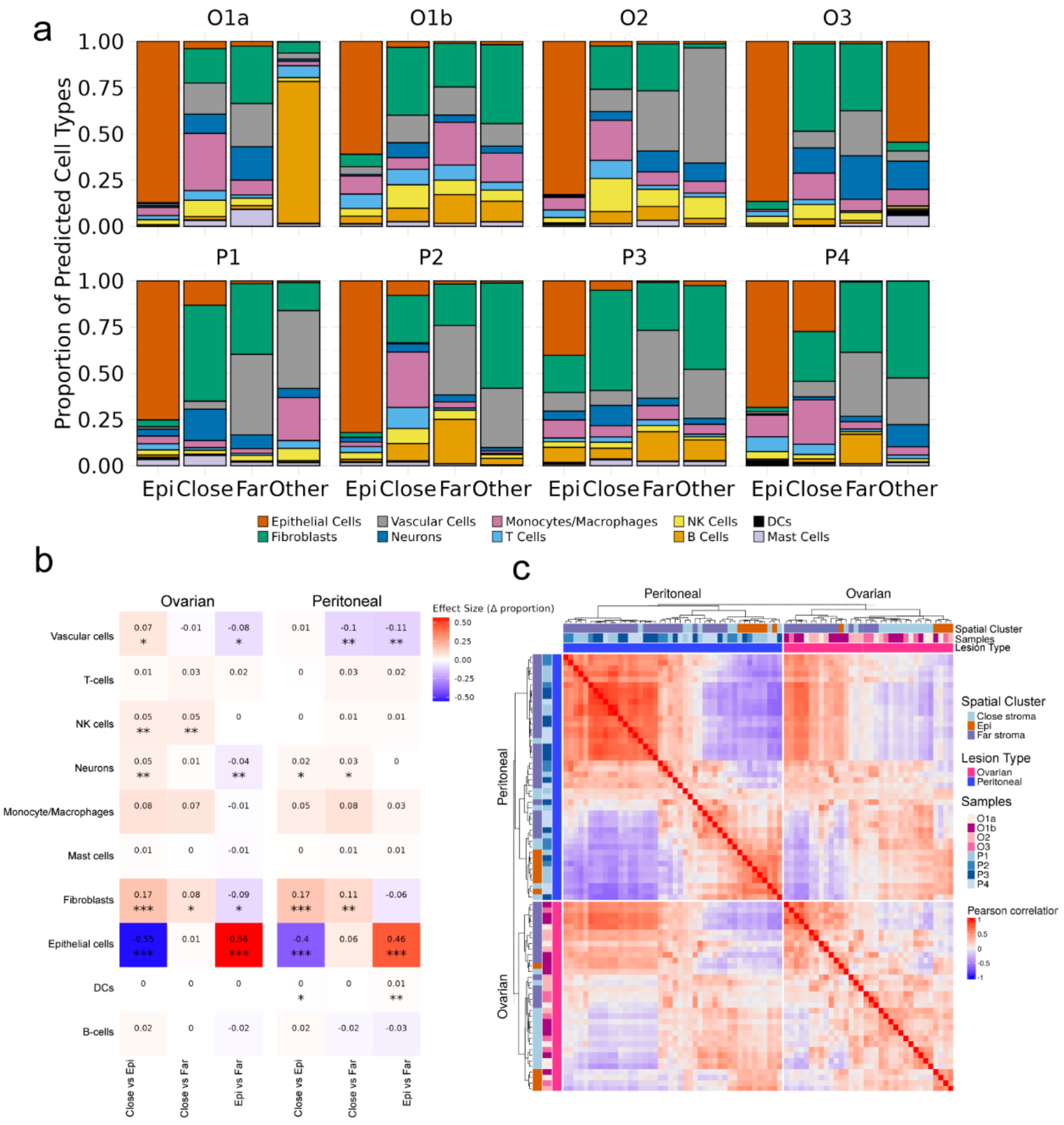


**Sup. Fig. 3: Predicted cell types, stats and supervised clustering of global cell types in endometriosis lesions.**

**a,** Proportion bar plot for each sample of ovarian lesions (top) and peritoneal lesions (bottom) of global cell types in each spatial cell cluster. **b,** Linear mixed-effect model heatmap of average cell type proportions in spatial cell clusters of ovarian and peritoneal lesions from Fig. 4c,d. The y-axis shows the cell types, and the x-axis shows the spatial cell cluster comparison. Number in each box is the effect size that correlates with the color scale, with red (positive effect) being increased in the first cluster compared and blue (negative effect) being higher in the second cluster compared. Stars represent significance: * $p < 0.05$, ** $p < 0.01$, *** $p < 0.005$. **c,** Heatmap of supervised clustering separating peritoneal and ovarian lesions comparing spatial cell clusters and samples.

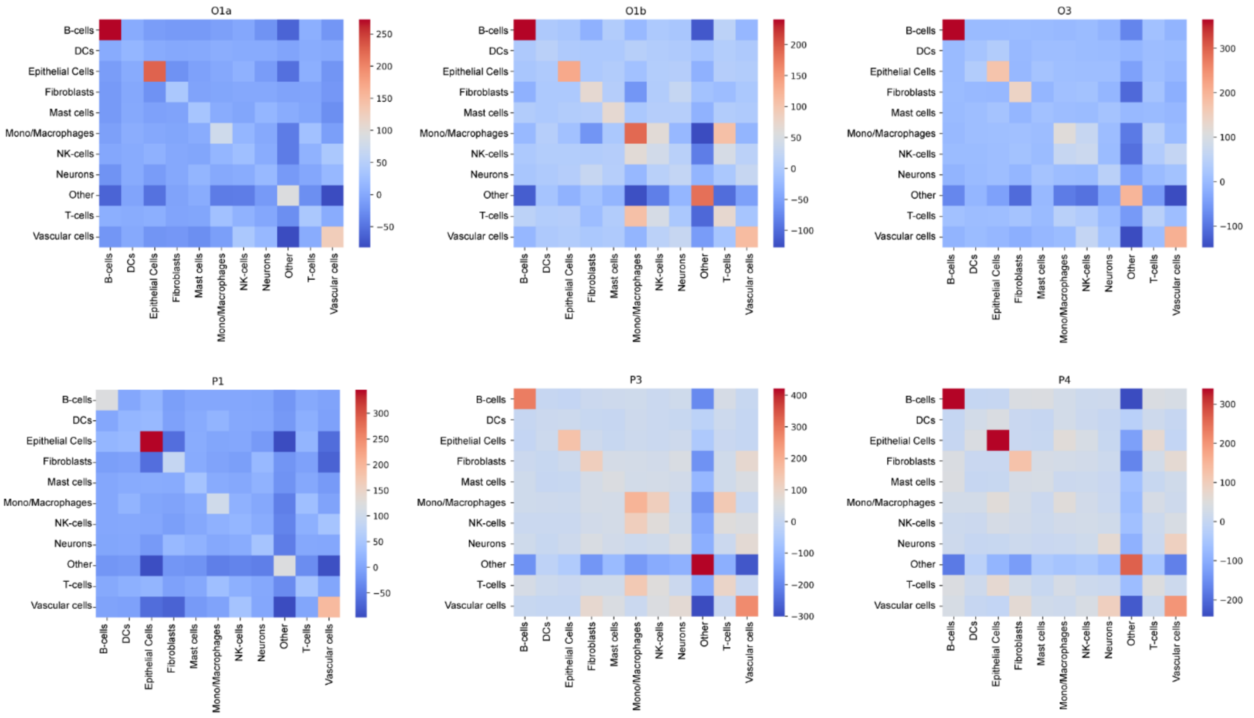


**Sup. Fig. 4: Neighborhood analysis of global cell types per sample.**
Neighborhood analysis comparing distances between global cell types per sample, ovarian lesions (top) and peritoneal lesions (bottom).

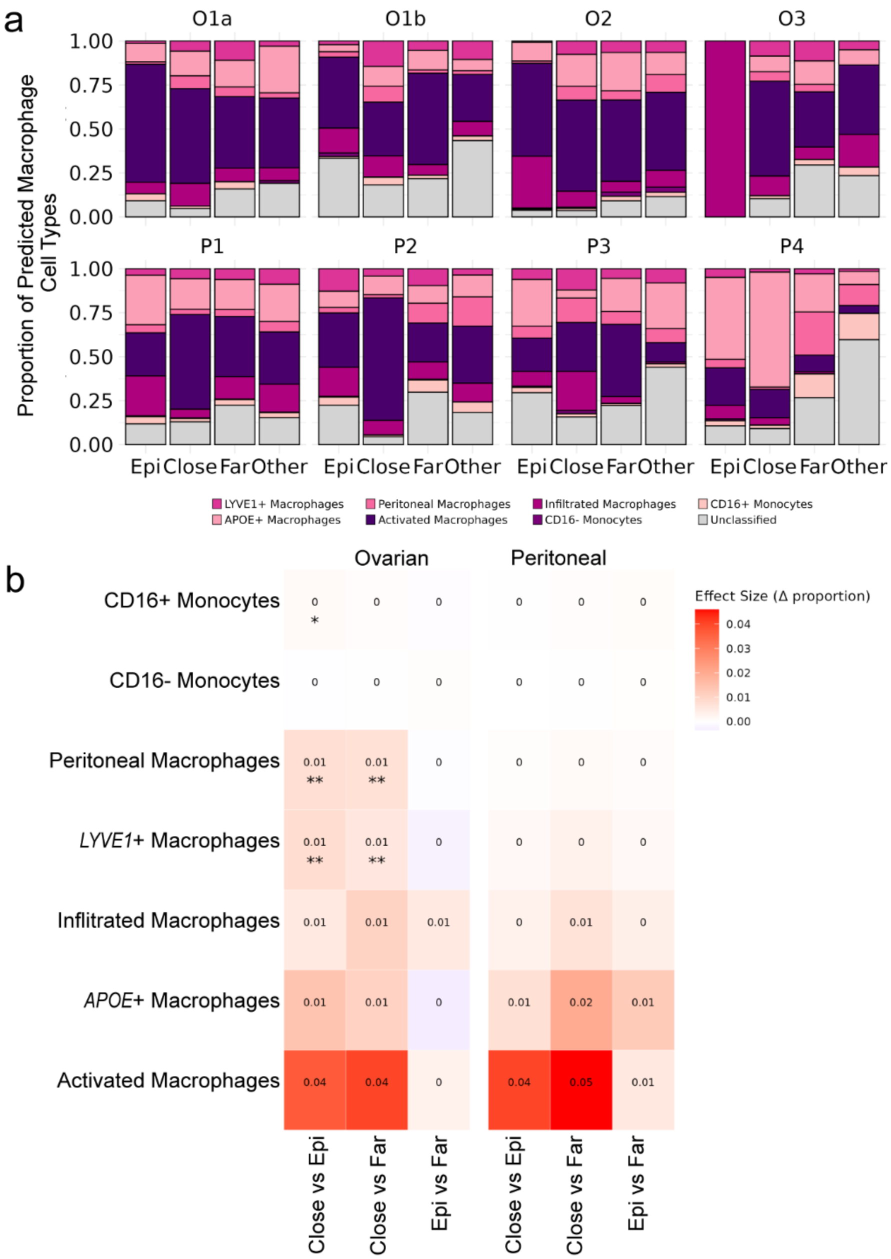


**Sup. Fig. 5: Per sample monocyte/macrophage subpopulation proportions and statistics.** **a,** Per sample monocyte/macrophage subpopulation proportions in ovarian lesions (top) and peritoneal lesions (bottom). **b,** Linear mixed-effect model heatmap of average monocyte/macrophage subpopulation proportions in spatial clusters of ovarian and peritoneal lesions from Fig. 5f. The y-axis shows the cell types, and the x-axis shows the spatial cluster comparison. Number in each box is the effect size that correlates with the color scale, with red (positive effect) being increased in the first cluster compared and blue (negative effect) being higher in the second cluster compared. Stars represent significance: * $p < 0.05$, ** $p < 0.01$, *** $p < 0.005$.

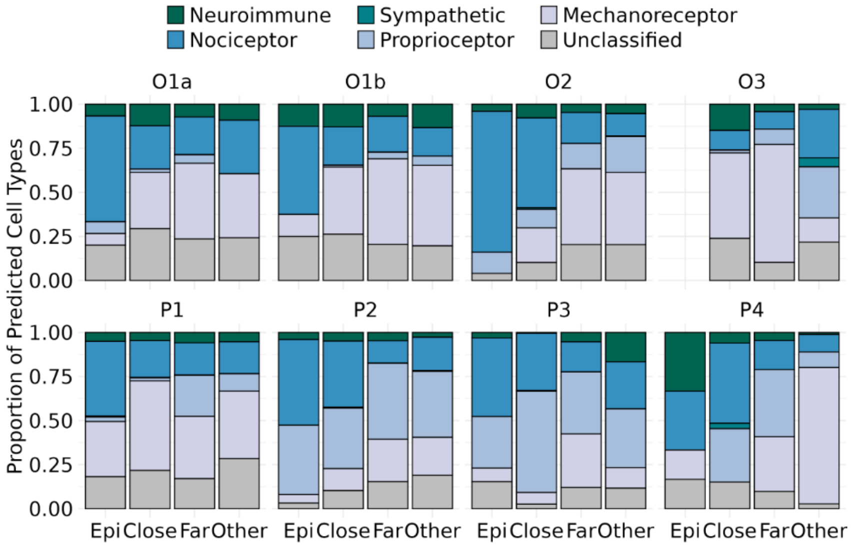


**Sup. Fig. 6: Per sample neuronal subtype proportions.**

Per sample neuronal subpopulation proportions in ovarian lesions (top) and peritoneal lesions (bottom).

a

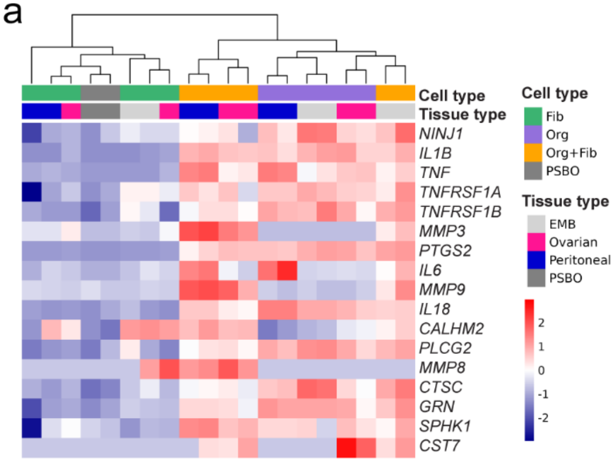


**Sup. Fig. 7: Neuro-immune related molecules upregulated in epithelial cell in vtiro co-culture conditions.**

**a,** Heat map of genes related to NINJ1, macrophage and inflammation signaling across the co-culture conditions described in Fig. 7a.

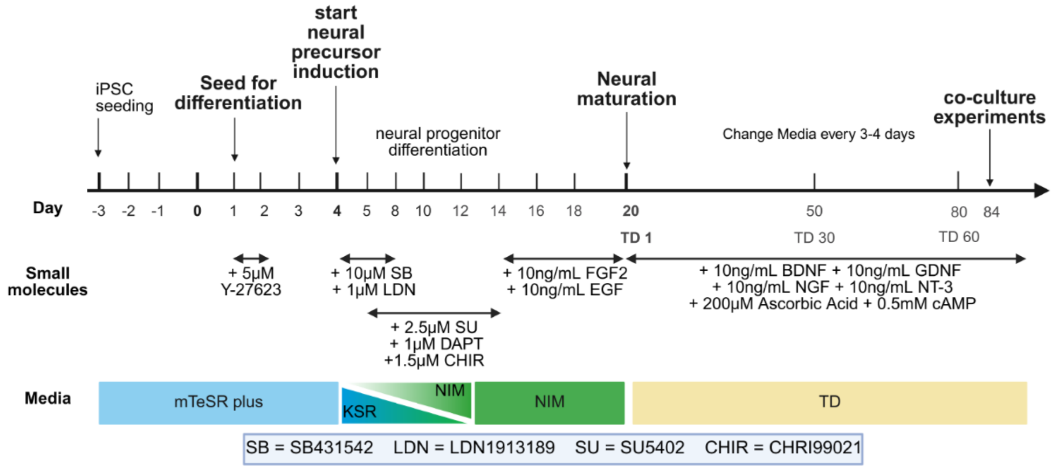


**Sup. Fig. 8: Differentiation protocol for iPSC derived peripheral sensory brain organoids.**
